# Target lipidomics reveals associations between serum sphingolipids and insulin sensitivity by the glucose clamp

**DOI:** 10.1101/2020.10.23.352765

**Authors:** Jingya Ye, Xuan Ye, Wanzi Jiang, Chenyan Lu, Xiaomei Geng, Chenxi Zhao, Yizhe Ma, Panpan Yang, Sin Man Lam, Guanghou Shui, Tao Yang, John Zhong Li, Yingyun Gong, Zhenzhen Fu, Hongwen Zhou

## Abstract

**Background:** This study aimed to systematically investigate the associations between serum sphingolipids and insulin sensitivity as well as insulin secretion. This study also aimed to reveal potential predictors for insulin sensitivity or give perceptive insight into disease processes.

**Methods:** We conducted a lipidomics evaluation of molecularly distinct SPs in the serum of 86 consecutive Chinese adults with or without obesity and diabetes using electrospray ionization mass spectrometry coupled with liquid chromatography. The GIR30 was measured under steady conditions to assess insulin sensitivity by the gold standard hyperinsulinemic-euglycemic clamp. We created the ROC curves to detect the serum SMs diagnostic value and establish the diagnosis of insulin sensitivity.

**Results:** Differential correlation network analysis illustrated correlations amongst lipids, insulin sensitivity, insulin secretion and other clinical indexes. Total and subspecies of serum SMs and globotriaosylceramides (Gb3s) were positively related to GIR30, free FAs (FFA 16:1, FFA20:4), some long chain GM3 and complex ceramide GluCers showed strong negative correlations with GIR30. Notably, ROC curves showed that SM/Cer and SM d18:0/26:0 may be good serum lipid predictors of diagnostic indicators of insulin sensitivity close to conventional clinical indexes such as 1/HOMA-IR (all areas under the curve >0.80) based on GIR30 as standard diagnostic criteria.

**Conclusions:** These results provide novel associations between serum sphingolipid between insulin sensitivity measured by the hyperinsulinemic-euglycemic clamp. We further identify two specific SPs that may represent prognostic biomarkers for insulin sensitivity.

## Background

SPs, including SMs, GM3s, ceramides, and gangliosides, are a group of ubiquitously produced lipids that play crucial roles in cell membrane function and cell signaling pathways(1). Sphingolipids and their substrates and constituents, FAs, have been implicated in the pathogenesis of various metabolic diseases associated with decreased insulin sensitivity/IR, obesity, diabetes, and atherosclerosis(2,3). When β cells of obese and IR individuals fail to secrete enough insulin to compensate for decreased insulin sensitivity, hyperglycemia develops(4). It is likely that perturbations in serum SPs and FAs are associated with diseases. Identifying these associations may reveal useful predictors or give perceptive insight into disease processes. For instance, decreased circulating saturated FAs through food intake, a kind of harmful FAs, could improve mouse insulin sensitivity(5). Cell membrane GM3s were found to display opposite associations with insulin resistance depending on acyl chain compositions(6). Based on a study of 2302 Singapore Chinese population, two specific SPs might represent prognostic biomarkers for T2DM (7). Another study found that SMs C14:0, C22:3, and C24:4 were positively associated with insulin secretion and glucose tolerance(8). However, due to the complexity and diversity of lipids, and differences in the analytical approaches adopted for sphingolipid measurements, the precise associations of sphingolipids with clinical indices of insulin sensitivity and secretion remain unclear. Fortunately, given the fundamentally development of LC-MS, the overall classification of lipid species in biological processes and has been improved in recent years. Such advances have expanded the collection of the “sphingolipidome” to more than 600 structurally distinct SPs and potentially thousands of theoretical metabolites that are likely to exist(9). In our study, we used a profile with more comprehensive coverage of different SP subclasses to recover their novel correlations with insulin sensitivity.

Commonly used clinical indexes for the evaluation of insulin sensitivity/resistance, such as the HOMA-IR, Matsuda index and QUICKI, are derived from circulating glucose and insulin levels, which are simple and noninvasive but indirect and subject to fluctuation (10). The gold standard for assessing insulin sensitivity is the hyperinsulinemic-euglycemic clamp developed by the DeFronzo group in the 1970s (11). This approach assesses insulin-mediated glucose utilization under steady-state conditions, which is both laborious and time-consuming as well as limited for large-scale use. However, the advantages of the glucose clamp are obvious for measuring insulin sensitivity/resistance directly and straightforwardly and further distinguishing hepatic and peripheral insulin sensitivity/resistance simultaneously when radiolabeled glucose tracers are used. To date, the glucose clamp has been applied as a standard assessment for other clinical indexes in various cross-sectional studies and prospective investigations to test the effect of different interventions on insulin sensitivity (12–14). Moreover, the IVGTT is an alternative method recommended to test insulin secretion and evaluate pancreatic islet function. It is acknowledged that a reduction in FPIR is the earliest detectable defect in β-cell function in individuals predisposed to develop T2DM and that this defect largely indicates β-cell exhaustion after years of compensation for antecedent IR (15).

SP metabolism is dysregulated in metabolic diseases and any of these specific lipid molecules may represent a potential biomarker of insulin resistance or other metabolic diseases. To date, there have been few systematic evaluations of the correlations between individual serum SP species with insulin sensitivity assessed by the hyperinsulinemic-euglycemic clamp. In this study, we investigated the associations of various serum SPs and FAs quantitated using a targeted lipidomic approach with insulin sensitivity and insulin secretion(Figure 1). We aimed to 1) investigate the incipient SPs and FAs patterns related to insulin sensitivity and insulin secretion, 2) identify predictive and diagnostic serum lipid indicators related to insulin sensitivity or islet function, thus preventing elevated risks of related symptoms and associated treatment expenditures, and. 3) explore and discuss potential lipid metabolic pathways associated with insulin sensitivity or islet function.

**Figure 1.**
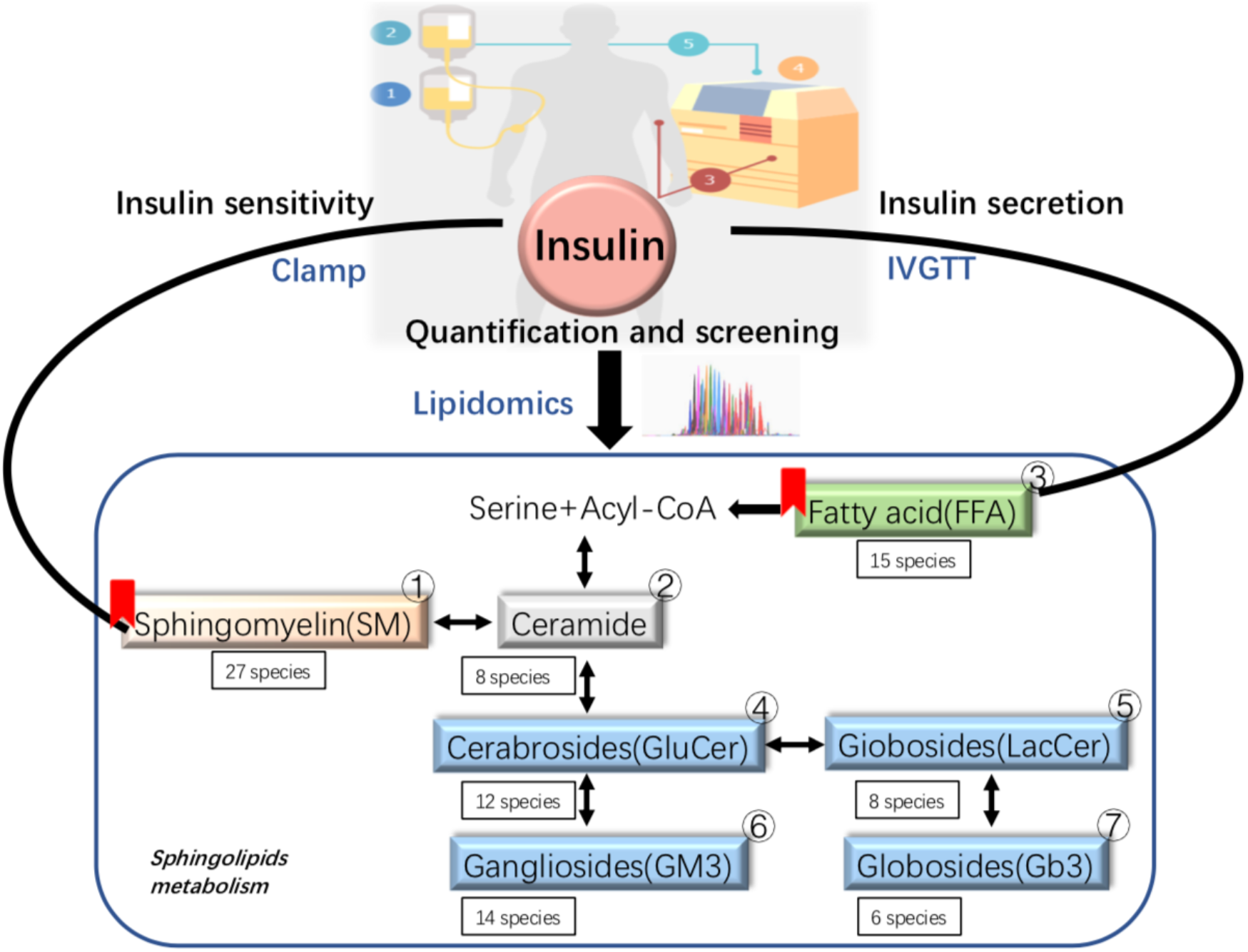
Study design. Decreased insulin sensitivity is a multifactorial condition related to obesity, increased inflammation and dyslipidemia. Lipotoxicity impairs islet function. Sphingolipids are a diverse class of lipids and we aimed to systematically investigate the changes in serum lipids in individuals and uncover potential serum sphingolipid predictors for insulin sensitivity and insulin secretion. Serum samples were analyzed by a targeted lipidomics approach. The glucose infusion rate over 30 minutes (GIR_30_) under steady conditions was measured by the gold standard hyperinsulinemic-euglycemic clamp, and first-phase insulin release (FPIR) was evaluated by the intravenous glucose tolerance test (IVGTT).

## Materials and Methods

### Study participants

Briefly, 86 participants aged from 18 to 60 years (number of Female:46) were enrolled in this project in Nanjing, Jiangsu Province, China. To broadly detect the range of insulin sensitivity, participants were normal weight (n=25), overweight(n=9; 24 kg/m2 ≤ BMI < 30 kg/m2) or obese(n=52(BMI > 30 kg/m^2^)), and with or without diabetes(number of participants with diabetes:25). Exclusion criteria included use of medication or supplements, smoking or high alcohol use (>4 and >2 standard drinks/week for men and women, respectively), other major diseases or presence of acute inflammation or pregnancy(Figure 2).

**Figure 2.**
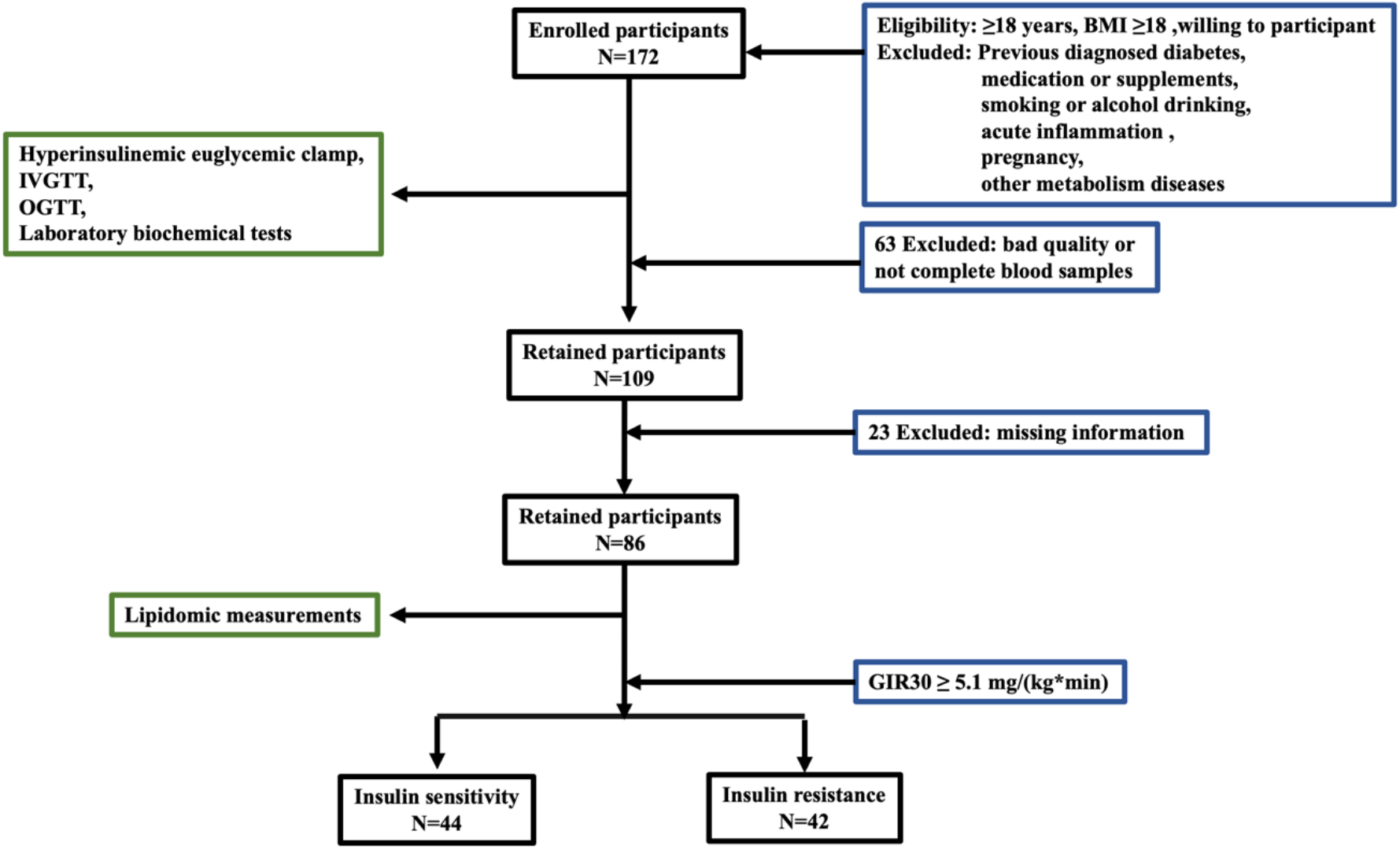
Flow chart of the study

### Hyperinsulinemic-euglycemic clamp

The hyperinsulinemic-euglycemic clamp presumes that the hyperinsulinemic state induced by a high dose of insulin infusion (>80 mU/m^2^*min) is sufficient to totally suppress hepatic glucose production and that there is no net change in the circulating blood glucose concentration under steady conditions(Supplementary Figure 1). Therefore, the GIR is equivalent to that of whole-body GDR or M value, which represents body insulin sensitivity(11). After an overnight fast, subjects were studied the next morning in the supine position. A prepared dose of insulin (80 mU/m^2^) was injected over the next 10 minutes to achieve a steady-state insulin concentration. The clamp duration was at least 2h. A variable infusion of 20% glucose was started to maintain the plasma glucose concentration at 5 mmmol/L (4.5-5.5 mmol/L). GIR_30_ was calculated from the mean glucose infusion rate during the last 30 minutes (14).

### IVGTT

Briefly, 0.3 g/kg body weight of a 20% glucose solution was given within 3 minutes. The FPIR is defined as the initial burst of insulin, which is released within the first 10 minutes and peaks at 2-4 minutes after glucose injection. After the acute response, there is a second-phase insulin secretion, which rises more gradually (16). Blood samples for the measurement of plasma glucose and serum insulin were collected at 0, 2, 4, 6, 8, 10, 20, 30, and 60 minutes. The FPIR was calculated as the mean of the 2- to 4-minute samples minus the mean pre-stimulus hormone concentration (0 minutes) (17).

### Laboratory measurements

Blood glucose was analyzed by a blood glucose biochemical analyzer (Germany, Biosen). Serum insulin and C-peptide were measured by chemiluminescent methods. HbA1c was analyzed by a chromatographic technique, and TC, TG, LDL-c, HDL-c and other biochemical phenotypes were measured with standard enzymatic assays in the laboratory of the First Affiliated Hospital of Nanjing Medical University, Nanjing, China.

### Targeted lipidomics analysis

Lipids were extracted from serum (20 μL) using a modified Bligh and Dyer’s extraction procedure (double rounds of extraction) and dried in a SpeedVac under OH mode. Prior to analysis, lipid extracts were resuspended in chloroform:methanol 1:1 (v/v) spiked with appropriate internal standards. All lipidomic analyses were carried out on an Exion UPLC system coupled with a QTRAP 6500 PLUS system (Sciex) as described previously(18). Sphingolipids were separated on a Phenomenex Luna Silica 3 μm column (i.d. 150×2.0 mm) under the following chromatographic conditions: mobile phase A (chloroform:methanol:ammonium hydroxide, 89.5:10:0.5) and mobile phase B (chloroform:methanol: ammonium hydroxide: water, 55:39:0.5:5.5) at a flow rate of 270 μL/min and column oven temperature at 25 °C. Individual sphingolipid species were quantified by reference to spiked internal standards including Cer d18:1/17:0, GluCer d18:1/8:0, LacCer d18:1/8:0, and SM d18:1/12:0, obtained from Avanti Polar Lipids; d_3_-GM_3_ d18:1/18:0 and Gb_3_ d18:1/17:0 purchased from Matreya LLC; d_8_-FFA 20:4 from Cayman Chemicals; and d_31_-FFA16:0 from Sigma-Aldrich.

### Statistical analysis

Lipids were divided into groups and analyzed based on carbon atom and double bond numbers. The typical cutoff value of GIR_30_ (or GDR, M value) of 4.9-5.2 mg/kg per minute was used to define insulin sensitive or resistant subjects(14,19). In our study, the cutoff point of GIR_30_ was set as 5.1 mg/(kg *min), which was also the median of consecutive 86 subjects. Differences in metabolic characteristics between two groups divided by GIR_30_ were calculated by t-test or the Mann-Whitney U test. Analyses were performed using SPSS for Macintosh version 25.0 (SPSS Inc, Chicago, IL, USA). Associations between metabolic parameters and lipid species were analyzed by Pearson correlation based on univariate logistic regression and adjusted for multiple factors, including age, sex, BMI, HDL-c, LDL-c, TG, TC, ALT and AST. Gephi was used to build correlation networks from differentially correlated lipid pairs. Data are presented as the mean ± SD or median (range). P<0.05 was considered statistically significant.

## Results

### Participants’ characteristics

Table 1 shows the characteristics of the 86 subjects in our study. According to the glucose clamp results, all the participants were divided into insulin-resistant and insulin-sensitive groups by GIR_30_, which was defined as 5.1 mg/(kg *min) (Table 1). We also applied IVGTT to evaluate islet function, and all individuals were separated into high insulin secretion and low insulin secretion groups by FPIR (Table 2). Subjects who presented insulin sensitivity were similar in age and sex to IR individuals but had lower levels of serum fasting glucose and 2h-OGTT glucose, neutral lipids including TC, TG, HDL-c, and LDL-c and liver enzymes including ALT and AST. Those who showed lower FPIR had comparable age, BMI and neutral lipids but higher levels of serum fasting glucose and 2h-OGTT glucose than those with high FPIR.

**Table 1.** Participants’ characteristics divided by GIR_30_ (mg / (kg *min))

|  | Insulin sensitivity (GIR<sub>30</sub> ≥ 5.1) |  | Insulin resistance (GIR<sub>30</sub> < 5.1) |  | P value |
| --- | --- | --- | --- | --- | --- |
|  | Mean ± SD | Median(range) | Mean ± SD | Median(range) |  |
| No.(F) | 44(25) |  | 42(21) |  | 0.53<sup>a</sup> |
| Age (years) | 27.6±8.1 | 25.0(18.0-53.0) | 29.4±8.5 | 28(18.0-56.0) | 0.08<sup>b</sup> |
| BMI (kg/m<sup>2</sup>) | 26.2±7.5 | 23.3(18-47) | 40.4±9.1 | 38.5(28.0-69.9) | <0.001<sup>b</sup> |
| FBG (mmol/L) | 5.1±0.6 | 4.9(4.2-7.1) | 6.8±2.7 | 5.5(4.3-17.3) | <0.001<sup>b</sup> |
| 2hPG (mmol/L) | 6.8±2.7 | 6.0(3.5-20) | 11.1±4.6 | 10.7(3.7-21.6) | <0.001<sup>c</sup> |
| TC (mmol/L) | 4.4±0.8 | 4.2(2.8-6.3) | 5.3±1.1 | 5.4(2.9-7.9) | <0.001<sup>c</sup> |
| TG (mmol/L) | 1.1±0.6 | 0.9(0.4-2.9) | 2.6±3.7 | 1.8(0.7-23.9) | <0.001<sup>b</sup> |
| HDL-c (mmol/L) | 1.4±0.3 | 1.3(0.8-2.1) | 1.1±0.2 | 1.1(0.7-1.6) | 0.01<sup>c</sup> |
| LDL-c (mmol/L) | 2.6±0.7 | 2.5(1.0-4.5) | 3.4±0.9 | 3.4(1.1-5.5) | <0.001<sup>b</sup> |
| ALT (U/L) | 32.7±33.0 | 18(8.0-160.6) | 84.5±79.9 | 68.4(6.1-404.8) | <0.001<sup>b</sup> |
| AST (U/L) | 28.3±20.1 | 20.5(13.0-104.7) | 58.0±56.1 | 43.2(11.2-319.0) | <0.001<sup>b</sup> |

F (female), body mass index (BMI), alanine aminotransferase (ALT), aspartate aminotransferase (AST), total cholesterol (TC), triglyceride (TG), high density lipoprotein cholesterol (HDL-c), low density lipoprotein cholesterol (LDL-c), fasting blood glucose (FBG), 2-hour plasma glucose (2hPG). Data are presented as the mean ± SD or median (range). <sup>a</sup> x<sup>2</sup> test. <sup>b</sup> Mann-Whitney U test. <sup>c</sup> Independent samples t test.

### Serum lipids and basic clinical characteristics

A. total of 90 lipid species in 7 classes of lipids, including FFAs, SMs, ceramides, GluCers, LacCers, GM3s and Gb3s, were quantitated and analyzed in the 86 human serum samples. We first evaluated differences among lipids with age and gender. There was a tendency of increased SMs and significant enhancements in GluCer d18:0/18:0 and Gb3 d18:1/20:0 (P<0.05) and decreases in GM3 d18:1/18:0, GM3 d18:0/18:0, and GM3 d18:1/20:0 (P<0.05) in females compared with males (P<0.05); however, no differences were found in the total or subtypes of serum ceramides (Supplementary Table 1). The lipid classes that extensively increased with age were FFAs, ceramides and GM3 (P<0.05), and those that tended to drop were SMs, GluCer and LacCer (Supplementary Table 2). Similarly, in individuals with high BMI values, the perceivably increased lipids were FFAs, ceramides and GM3s, while lipids such as SMs, GluCers and LacCers declined (Supplementary Table 3).

### Differential analysis and correlation network of GIR and FPIR

We used a heatmap to visualize the trends of lipidomics among the 86 individuals divided by GIR_30_ (Figure 3, Supplementary Table 4). The results showed a significant higher in total serum SMs in the high GIR_30_ (insulin sensitive) group (P<0.05) and a mild tendency of serum GluCers, LacCers and Gb3s, especially GluCer d18:1/20:0 and Gb3 d18:1/24:0, to increase. However, in the high GIR_30_ group, a decrease was also seen in some FFAs, ceramides and GM3s (Figure 3), especially C18 FFAs (FFA 18:3, FFA 18:1, FFA 18:0) and FFA16:1, long-chain unsaturated ceramides (Cer d18:1/22:0, Cer d18:1/24:0, Cer d18:1/24:1) and most saturated GM3s (GM3 d18:0/16:0, GM3 d18:1/18:0, GM3 d18:0/18:0, GM3 d18:1/20:0,GM3 d18:0/20:0, GM3 d18:1/22:0, GM3 d18:0/22:0, GM3 d18:0/24:0). Interestingly, most serum lipids increased in individuals with low insulin secretion, including FFAs (FFA 22:6, FFA 22:5, FFA 18:2, FFA 18:1, FFA 18:0, FFA 16:0), Lac-Cers (LacCer d18:1/16:0, LacCer d18:1/18:0, LacCer d18:1/20:0, LacCer d18:1/22:0, LacCer d18:1/24:0, LacCer d18:1/24:1), ceramide (Cer d18:1/22:0, Cer d18:1/24:0, Cer d18:1/24:1), and GM3 (GM3 d18:0/16:0, GM3 d18:1/18:0, GM3 d18:0/20:0, GM3 d18:0/24:0, GM3 d18:1/22:0, GM3 d18:0/22:0, GM3 d18:1/24:0, GM3 d18:0/24:0) (Supplementary Table 5). These findings might be because of lipid toxicity.

**Figure 3.**
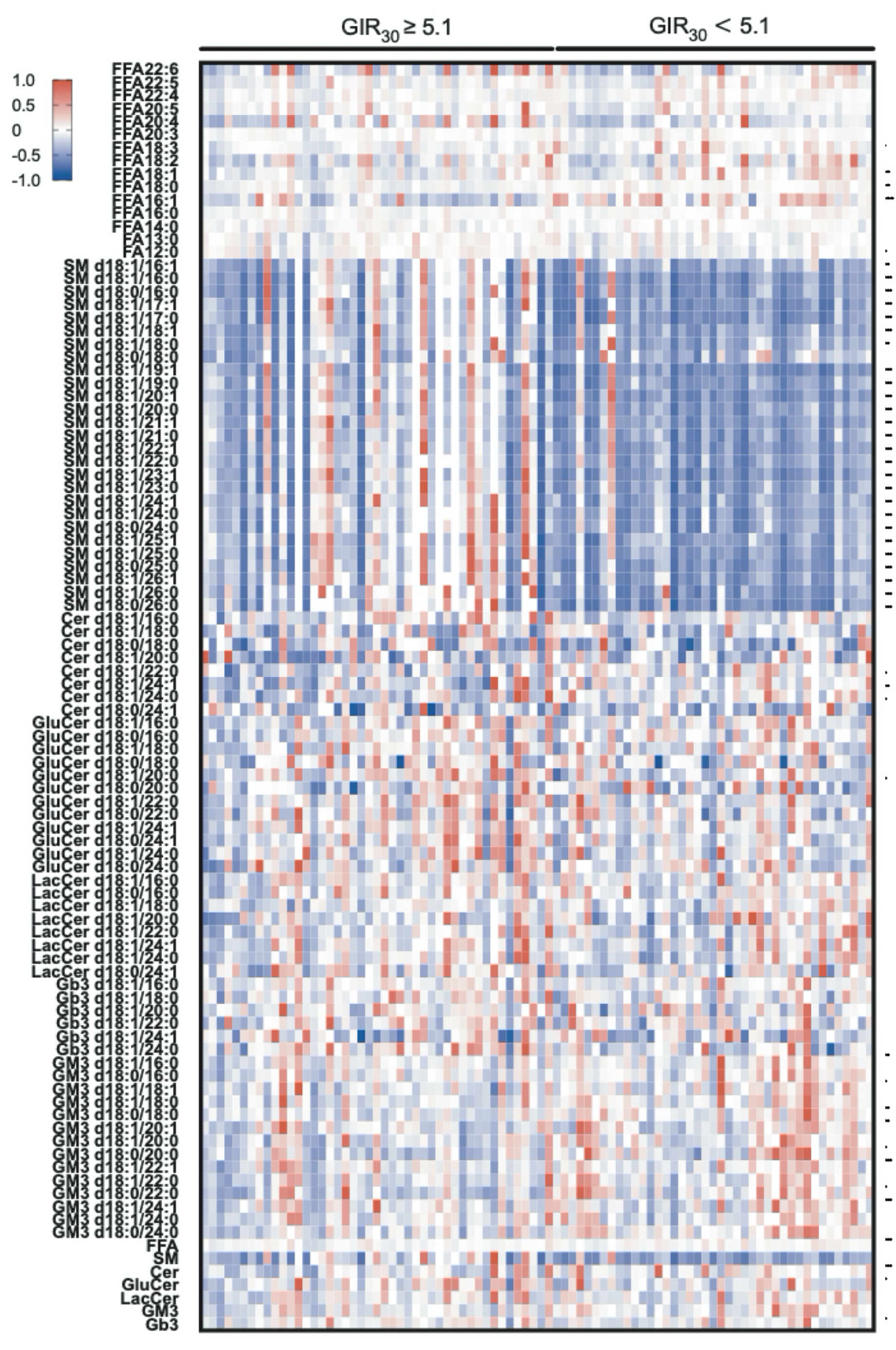
Serum lipid concentrations in individuals visualized by heatmap. Each colored cell on the map corresponds to a concentration value. The concentration values were transformed to −1 (most below average, Blue) to 1 (most above average, Red). The 90 lipid species from the two groups are presented and range from insulin sensitivity to IR. s: non-IR group, r IR group.

A correlation network was constructed to systematically evaluate the correlation of the clinical indexes and lipidomics, as well as lipid-related interrelationships. We raised our correlation coefficient limit to 0.2 (p<0.05), and the results indicated that the lipids that were positively related to GIR_30_ including GluCers (GluCer d18:1/20:0, GluCer d18:1/22:0, GluCer d18:0/24:0, GluCer d18:0/24:1), Gb3s (GB3 d18:1/24:0, GB3 d18:1/16:0) and all SMs (SM d18:1/19:1, SM d18:1/23:1, SM d18:1/25:1, SM d18:1/25:0, SM d18:0/25:0, SM d18:1/26:1, SM d18:1/26:1, SM d18:0/26:0, and so on), while FFAs (FFA16:1, FFA20:4) and GM3 (GM3 d18:1/18:0, GM3 d18:0/18:0, GM3 d18:1/20:0, GM3 d18:0/20:0) were negatively correlated with GIR_30_. This relationship still existed when adjusted for age, sex, BMI, TG, TC, HDL-c, LDL-c, ALT and AST (Figure 4 Supplementary Table 6). Serum lipids showed a negative relationship with FPIR, including FFAs (FFA22:5, FFA18:2, FFA18:1, FFA16:0), GluCers (GluCer d18:1/16:0, GluCer d18:1/18:0, GluCer d18:1/20:0), LacCer (LacCer d18:1/20:0, LacCer d18:1/24:0, LacCer d18:1/24:1), which still existed after adjustment for age, sex and BMI (Figure 4, Supplementary Table 6). Notably, HDL is a lipid protein known to transport SMs. Our data showed that HDL-c, not LDL-c, was correlated positively with most SMs.

**Figure 4.**
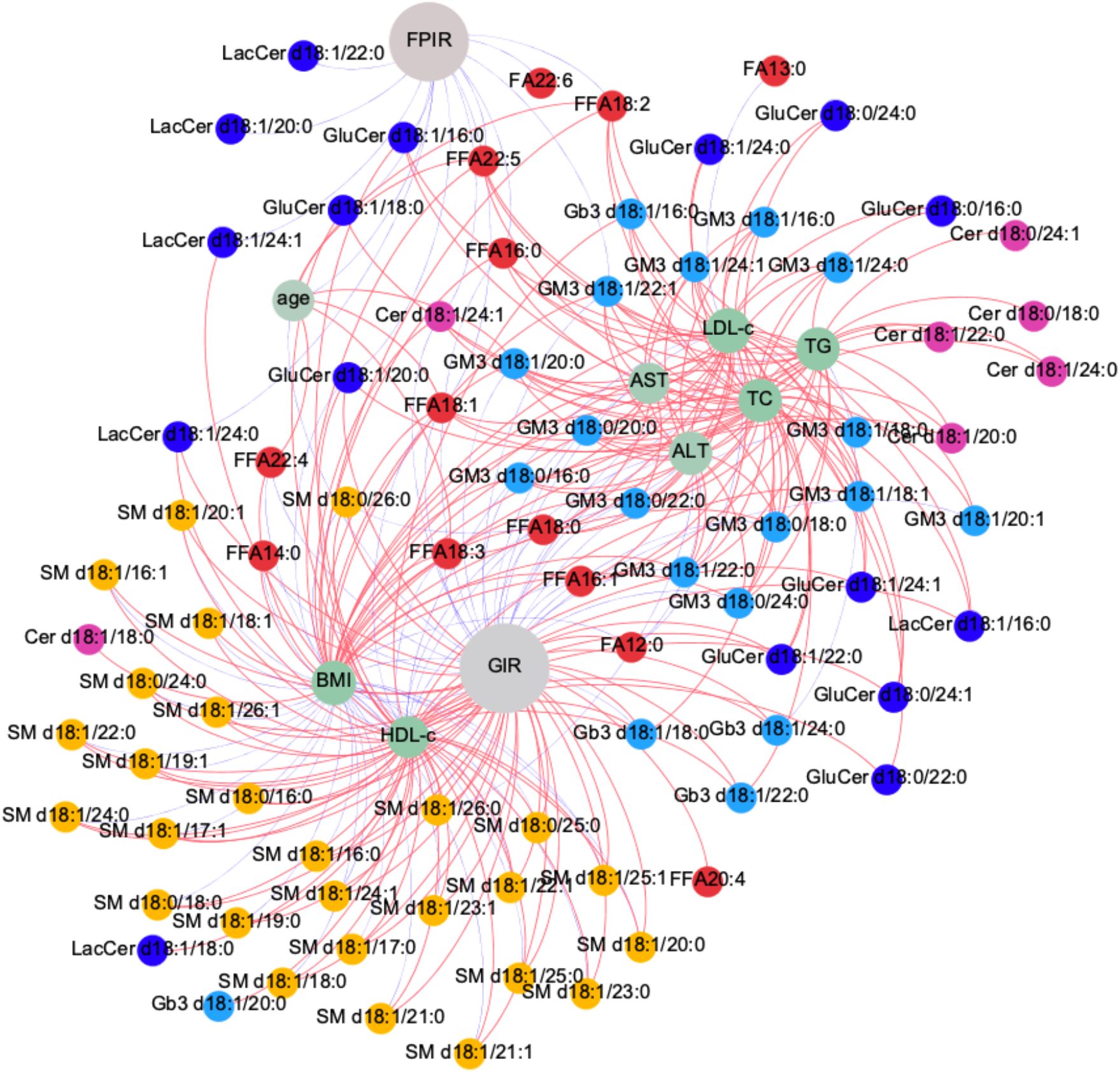
Correlations between serum lipids and multiple clinical characteristics. Clinical characteristics network showing lipid correlations with GIR, FPIR (grey nodes) and 8 key clinical characteristics (green nodes). Pearson’s correlation was analyzed (A) and absolute coefficient values of 0.2 and above are shown (P<0.05). Red edge color indicates positive correlation, while blue edge indicates negative correlation. Node color is grouped according to lipid type. SM nodes are yellow, ceramide nodes are pink, GluCer and LacCer nodes are dark blue, FFA nodes are red, and GM3 and Gb3 nodes are blue.

### Use of the ROC test for the diagnosis of insulin sensitivity

To discover serum lipid predictive and prognostic biomarkers for insulin sensitivity evaluation, we assessed the predictive value of various lipids by ROC curve analysis using GIR as a standard criterion (Figure 5, Supplementary Table 7). Notably, the ratio of total serum SMs and ceramides (SM/Cer) showed high diagnostic efficiency, as it was equal to 1/HOMA-IR, with 0.85 (0.77-0.94) and 0.85 (0.77-0.93), respectively. Moreover, SM 18:0/26:0, a subspecies of SMs, showed close and high diagnostic efficiency with 0.84 (0.75-0.93), while other lipids showed lower diagnostic efficiency, such as total serum ceramides 0.35 (0.23-0.46) and FFAs 0.38 (0.26-0.5).

**Figure 5.**
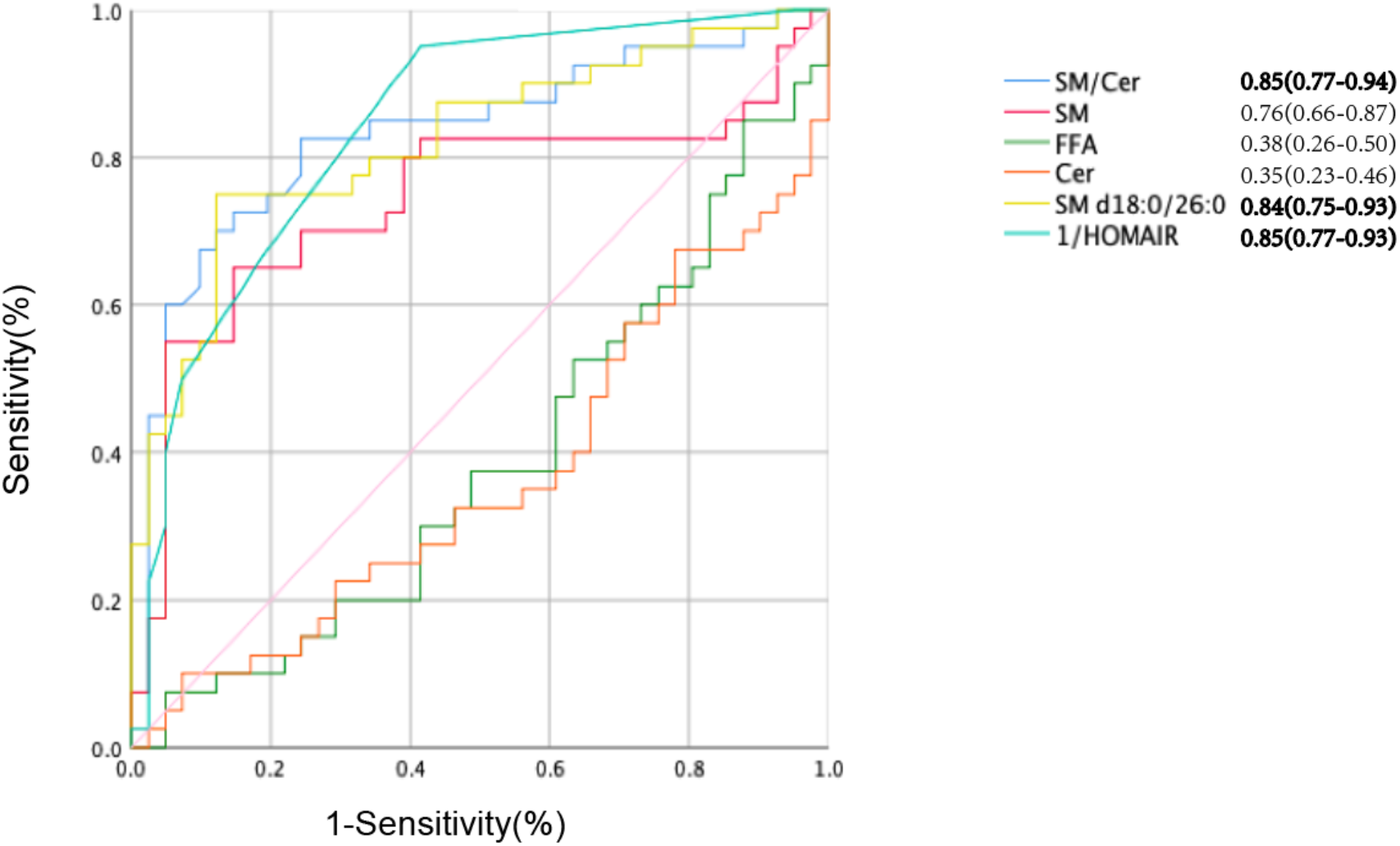
Different noninvasive clinical models or serum biomarkers for defining IR. Plots of area under the curves (AUC) in the total subjects using GIR ≥ 5.1 mg / (kg *min) as a reference. SM, total serum sphingomyelins; FFA, total serum fatty acids; Cer, total serum ceramides; SM/Cer, total serum sphingomyelins/ total serum ceramides. Model 1: 1/HOMAIR, AUC 0.852; Model 2: SM/Ceramide, AUC 0.851; Model 3, SM d18:0/26:0, AUC 0.839.

## Discussion

In the current work, we report the systematic analysis of the sphingolipidome, allowing for an accurate evaluation of the associations between serum SPs content and obesity and diabetes related characteristics, such as insulin sensitivity, BMI, blood lipids, and HOMA-IR. We also found that higher concentration of serum free FAs(FFA 22:6, FFA 22:5, FFA 18:2, FFA 18:1), substrates and constituents of SPs, were associated with a higher risk of decreased insulin secretion. Of note, ROC curves showed that SM/Cer and SM d18:0/26:0 may be good serum lipid predictors of diagnostic indicators for insulin sensitivity close to conventional clinical indexes such as 1/HOMA-IR (all areas under the curve >0.80) based on GIR_30_ as standard diagnostic criteria.

We conducted a lipidomics evaluation of molecularly distinct SPs in the serum of 86 consecutive Chinese adults with or without obesity and diabetes using electrospray ionization mass spectrometry coupled with liquid chromatography. Our study involved a thorough evaluation of lipid profile changes in human serum samples among 90 lipid species. These lipids are from 7 subclasses, including FFAs (15 species), SMs (27 species), ceramides (8 species), GluCers (12 species), LacCers (8 species), GM3s (14 species) and Gb3s (6 species). We found an increasing tendency in complex serum sphingolipids (SPs), including SMs and GluCers, in females, which is consistent with recent large-scale lipidomic investigations of Chinese T2DM patients in Singapore (7). The possible reason for sex differences might be sex hormone estrogens, which could improve sphingolipid synthesis and reduce sphingolipid degradation by attenuating acid sphingomyelinase activities (20,21). We also observed that serum FFAs, ceramides and GM3 were significantly higher in individuals of older age or with a higher BMI, which might be due to the exacerbation of lipid-related inflammatory factors with age or BMI (7,22–24). Interestingly, our results showed that SMs showed no significant changes with age or BMI, indicating that SMs were less likely to be age- or BMI-related proinflammatory mediators.

The glucose infusion rate over 30 minutes (GIR_30_) under steady conditions to assess insulin sensitivity by the gold standard hyperinsulinemic-euglycemic clamp, and first-phase insulin release (FPIR) to evaluate insulin secretion by the intravenous glucose tolerance test (IVGTT).

In comparison to previous studies (7,25,26), we further identified more lipid species related to obesity and T2DM. In general, serum FFAs, ceramides and GM3s were decreased in subjects with high GIR_30_; however, serum SMs, GluCers, LacCers and Gb3s were increased in individuals with high GIR_30_. Remarkably, all species of serum SMs analyzed were relevant to increased insulin sensitivity, while partial species of serum ceramides (Cer d18:1/22:0, Cer d18:1/24:0, Cer d18:1/24:1) were corroborated with a higher risk of impaired insulin sensitivity. In mammals, substrate specificity is distinguished by CerS with N-acyl chain length. CerS has six isoforms (CerS1-6), and CerS2 seems to be a unique enzyme with a higher activity for very long-chain fatty acids such as C22-C26. Our findings showed that these very long-chain ceramides (Cerd18:1/22:0, Cerd18:1/24:0, Cer d18:1/24:1) were increased in IR individuals, consistent with part of a previous study indicating that ceramides with larger carbon atom numbers and more unsaturated bonds were more strongly associated with an increased risk of diabetes (7). These findings suggested that compositional variations in specific species of ceramides were associated with risks of IR or diabetes, even though the number of total ceramides remained unchanged. We also found a positive correlation between serum GluCers and GIR_30_, and the paradox of correlations indicated that more complex ceramides are different from ceramide regulation. Cross-sectional studies about serum SMs and insulin sensitivity in humans are limited. Among these studies, the associations between SMs and HOMA-IR are still controversial (25,27–29). This discrepancy may be attributed to a number of factors: 1) ethnic characteristics, 2) effective population size, 3) age span changes from young adults to elderly individuals, 4) accuracy of clinical indicators such as HOMA-IR, Matsuda index or QUICKI, 5) different lipidomics analytical platforms, 6) combined influence of body weight, 7) onset of diabetes time course, 8) drug intervention for diabetes, and 9) accompaniment by other metabolic diseases. Interestingly, our results were notable: all species of SMs showed a strong positive relationship with GIR_30_, and only some ceramides were negatively related to GIR_30_. The characteristics of the study were as follows: 1) young and middle-aged adults between 18 and 56 (median 26.5), 2) body weight ranging from normal weight to overweight to obese, 3) obese subjects with or without newly diagnosed diabetes, 4) a thorough evaluation of SPs specific for 27 species of SMs and 8 species of ceramides through targeted lipidomics measurements, and 5) insulin sensitivity evaluated by the gold standard, glucose clamp evaluation.

We created the ROC curves to detect the serum SMs diagnostic value and establish the diagnosis of insulin sensitivity based on GIR_30_ derived from the glucose clamp, which is the standard method, and the cutoff point of GIR_30_ was set as 5.1 mg/(kg *min). ROC analysis provides tools to screen possibly optimal models in a direct and natural way of diagnostic decision making. It is a comparison of two operating characteristics as the criterion changes. In the current study, we found that serum very long-chain SM d18:0/26:0 and the ratio of total serum SMs and Cers (SM/Cer) could be good predictors of insulin sensitivity that were very close to 1/HOMA-IR. Although SM d18:0/26:0 accounted for the lowest portion of total serum SMs, it seems to be a much more sensitive indicator of insulin sensitivity. Ceramides are substrates that generate SMs and are intermediate lipid links in sphingolipid metabolism. We used SM/Cer as a novel index that reflects the direction of sphingolipid reactions. As the ratio increases, more SMs are produced in the forward direction. We aimed to discover the pathological relevance of our identified lipids with regard to insulin sensitivity. Because sphingolipid synthesis and catabolism are dysregulated in metabolic diseases, any of these specific molecules may reflect a potential biomarker of decreased insulin sensitivity or other pathologies (30). A study also revealed that a metabolite panel appreciably improved T2DM risk prediction over or close to conventional clinical factors (18,31). The serum lipid parameters we identified could be good predictors compared with HOMA-IR, where GIR_30_ was used as a standard reference. These predictors we screened emphasized that the scarce and uncommon lipids might be useful for further insulin sensitivity mechanism exploration as well as clinical diagnosis and prevention.

We did not find specific lipid indicators of insulin secretion by ROC analysis. However, our correlation network analysis data showed a negative association between serum total FFAs and FPIR, and it disappeared after adjustment for age, sex, BMI, TC, TG, HDL-c, LDL-c, ALT and AST. More interestingly, 18-chain FFAs and 22-chain FFAs remained negatively correlated with FPIR when adjusted for the above factors, which indicated a strong relationship between these subtypes of FFAs and FPIR. A previous study based on the OGTT observed an association between higher total fasting plasma FFA levels and lower insulin secretion in children and adults without adjustments for confounding factors (32). Some studies proved that chronic FFA-induced lipotoxicity on pancreatic islets contributed to β cell dysfunction (33,34). Therefore, a reduction in elevated plasma FFA levels might be a critical therapeutic target for obesity-related type 2 diabetes. Although some studies showed that 24-72 h fatty acid stimulation increased insulin secretion in isolated islets from nondiabetic mice or humans (35,36), there were two different notable effects of short-term and long-term FFA stimuli on insulin secretion, and β cell function was impaired by long-term FFA stimulation. We considered that the negative association between these subtypes of serum FFAs, including 18-chain and 22-chain FFAs and FPIR in our study population, represented long-term damage to insulin secretion.

The major strengths of our current study are as follows. First, our targeted lipidomics approach permitted the unambiguous identification and accurate quantitation of serum lipids. This procedure unveiled unambiguous identification and accurate quantitation of serum lipids and evaluated systematic lipid pathway coregulation via multiple correlation analyses. Second, in terms of the study cohorts, we screened serum lipid predictors for the diagnosis of insulin sensitivity in Chinese individuals including normal-weight, overweight, obese and diabetic subjects, which is the gold standard for evaluating insulin sensitivity clinically. Third, we found that serum very long-chain SM (d18:0/26:0) and the SM/Cer could be good predictors for insulin sensitivity, which was very close to 1/HOMA-IR and of great value for diagnosis and prevention. However, there were also some limitations. First, lifestyle habits, including diet and exercise, were not controlled in the current study due to the lack of formal questionnaire or recording. These habits may also influence the analysis of lipids, since some sphingolipid species were shown to increase in HFD-induced obese mice (37,38). Second, laborious and time-consuming glucose clamps limited our cross-sectional study from being performed on a large scale, and further work is required to verify our findings. Third, it might influence the serum sphingolipid patterns that IR patients in our study had higher BMI than those of insulin sensitivity, since we corrected these effects through statistics methods.

## Conclusion

In summary, these results provide novel associations between serum sphingolipid levels and insulin sensitivity measured by hyperinsulinemic-euglycemic clamp. This finding suggests that the balance of SMs metabolism, rather than ceramide, is correlated with the pathology of insulin resistance, obesity and T2DM. We further identified two specific SPs that may represent prognostic biomarkers for insulin sensitivity.

## Abbreviations

SP: sphingolipid
SM: sphingomyelin
GM3: sphingosine
FAs: fatty acids
GluCers: Glu-ceramides
LacCers: Lac-ceramides
GIR_30_: glucose infusion rate over 30 minutes
FPIR: first-phase insulin release
IVGTT: intravenous glucose tolerance test
ROC: receiver operating characteristic
IR: insulin resistance
LC-MS: liquid chromatography-mass spectrometry
QUICKI: quantitative insulin sensitivity check index
GDR: glucose disposal
M: metabolizable glucose
TC: total cholesterol
TG: triglyceride
LDL-c: LDL-cholesterol
HDL-c: HDL-cholesterol
ALT: alanine transaminase
AST: aspartate transaminase
Gb3: globotriaosylceramides
CerS: ceramide synthase
SM/Cer: ratio of total serum SMs and Cers.

## Ethics approval and consent to participate

This study protocol was approved by the Institutional Review Board of the First Affiliated Hospital of Nanjing Medical University, and all participants provided written informed consent (2014-SR-003). All participants provided written informed consent prior to study entry.

## Consent for publication

All authors have participated in the work and have reviewed and agree with the content of the article.

## Availability of data and materials

We do not have governance permissions to share individual level data on which these analyses were conducted since they derive from clinical record data. However, for any bona fide requests to audit the validity of the analyses, the verifiable research pipeline which we operate means that one can request to view the analyses being run and the same tabulations resulting. We are also happy to share summary statistics for those wishing to conduct meta-analyses with other studies.

## Authors’ contributions

The authors thank the staff and participants for their important contributions. Jingya Ye, Xuan Ye and Wanzi Jiang were responsible for data collection, analysis and writing of the first draft of manuscript. Zhenzhen Fu and Yingyun Gong contributed to the human glucose clamp study performance and guidance of clinical data acquisition. Jingya Ye, Xuan Ye, Wanzi Jiang, Xiaomei Geng, Chenxi Zhao, Chenyan Lu, Yizhe Ma, and Panpan Yang took part in human glucose clamp performance. Sin Man Lam and Guanghou Shui contributed to lipidomic measurement. John Zhong Li, Feng Chen and Tao Yang contributed to data analysis and statistic process. Hongwen Zhou was the guarantors of this work and, as such, had full access to all of the data in the study and take responsibility for the integrity of the data and the accuracy of the data analysis.

## Acknowledgements

The authors acknowledge all the physicians and nurses taking part in the enrolment of participants in the First Affiliated Hospital of Nanjing Medical University.

## Supplemental Material

**Supplementary Figure 1.**
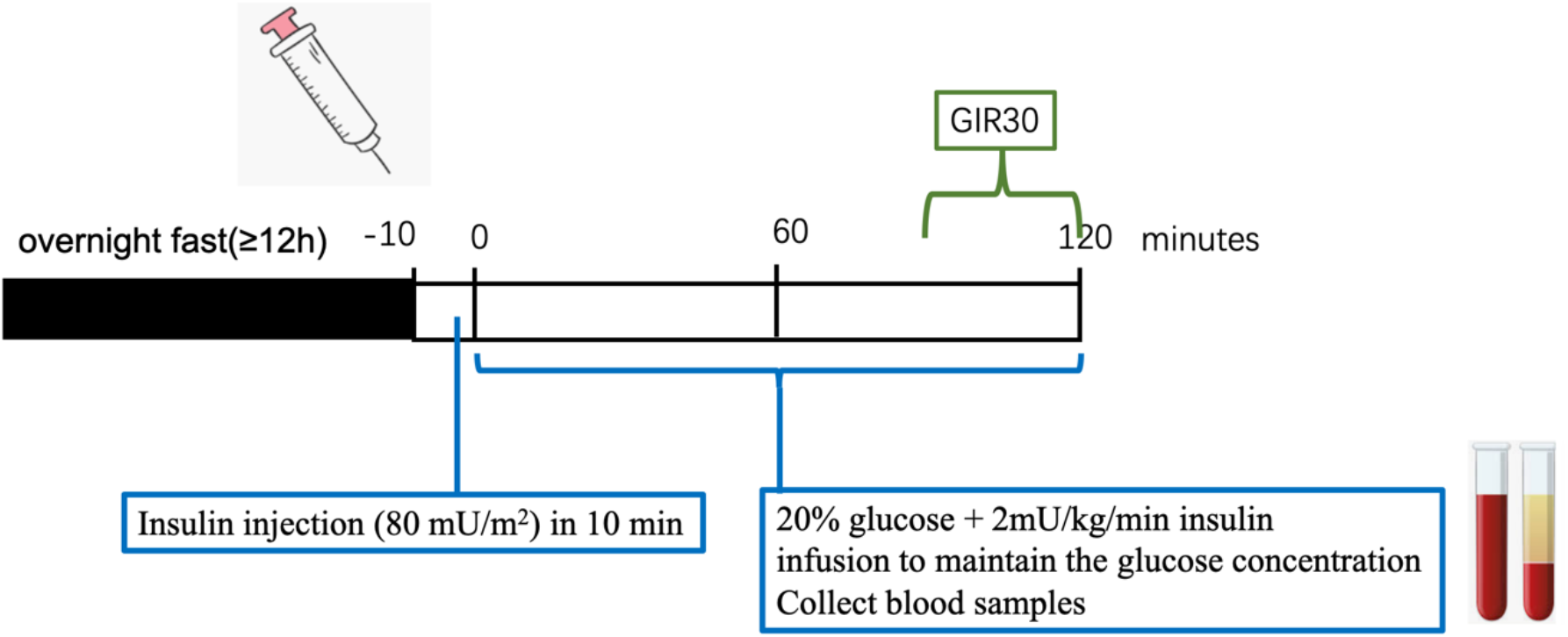
Flow chart of the hyperinsulinemic euglycemic clamp.

**Supplementary Table 1.** Gender differences in individual lipid species

| Lipid | Male (n=40) |  | Female(n=46) |  | P value |  | Percentage changed in Female /Male |
| --- | --- | --- | --- | --- | --- | --- | --- |
|  | Average lipid concentration (nM) | Standard deviation | Average lipid concentration (nM) | Standard deviation | t.test | Man whitney u test |  |
| FFA22:6 | 27,383.80 | 9,766.63 | 28,364.37 | 11,811.23 | 0.679 | 0.91 | 4% |
| FFA22:5 | 35,394.50 | 5,727.76 | 34,595.93 | 5,745.34 | 0.521 | 0.391 | -2% |
| FFA22:4 | 46,630.81 | 5,410.97 | 45,155.79 | 4,303.44 | 0.163 | 0.307 | -3% |
| FFA20:5 | 38,193.91 | 7,044.16 | 37,642.25 | 6,542.37 | 0.708 | 0.516 | -1% |
| FFA20:4 | 70,539.58 | 25,512.53 | 72,809.47 | 34,369.68 | 0.732 | 0.511 | 3% |
| FFA20:3 | 72,926.12 | 7,350.62 | 71,428.02 | 5,230.23 | 0.275 | 0.489 | -2% |
| FFA18:3 | 21,382.20 | 4,770.98 | 20,105.87 | 2,522.77 | 0.135 | 0.345 | -6% |
| FFA18:2 | 833,119.63 | 289,535.66 | 757,961.14 | 216,305.49 | 0.173 | 0.287 | -9% |
| FFA18:1 | 368,217.17 | 80,349.19 | 353,921.66 | 64,141.91 | 0.362 | 0.494 | -4% |
| FFA18:0 | 2,444,736.12 | 204,479.00 | 2,492,478.17 | 245,099.65 | 0.334 | 0.319 | 2% |
| FFA16:1 | 898,803.53 | 283,053.74 | 868,390.86 | 265,350.32 | 0.609 | 0.71 | -3% |
| FFA16:0 | 2,643,907.12 | 201,659.98 | 2,667,694.84 | 222,784.21 | 0.607 | 0.505 | 1% |
| FFA14:0 | 231,903.36 | 21,691.50 | 228,117.15 | 19,882.60 | 0.401 | 0.446 | -2% |
| FA13:0 | 1,393,717.86 | 189,564.71 | 1,381,101.56 | 165,049.25 | 0.742 | 0.368 | -1% |
| FA12:0 | 1,573,313.86 | 215,239.69 | 1,563,687.31 | 209,107.91 | 0.834 | 0.337 | -1% |
| SM d18:1/16:1 | 30,057.81 | 24,607.53 | 47,297.16 | 68,494.93 | 0.117 | 0.239 | 57% |
| SM d18:1/16:0 | 261,682.84 | 304,651.82 | 424,500.72 | 774,554.29 | 0.194 | 0.382 | 62% |
| SM d18:0/16:0 | 26,831.28 | 30,257.85 | 44,211.40 | 81,235.64 | 0.183 | 0.319 | 65% |
| SM d18:1/17:1 | 726.56 | 905.01 | 1,177.57 | 1,974.57 | 0.169 | 0.242 | 62% |
| SM d18:1/17:0 | 5,109.41 | 6,623.18 | 7,895.05 | 14,079.59 | 0.255 | 0.499 | 55% |
| SM d18:1/18:1 | 23,088.56 | 24,590.08 | 38,873.03 | 62,262.07 | 0.119 | 0.069 | 68% |
| SM d18:1/18:0 | 50,774.22 | 50,941.15 | 79,784.17 | 127,935.06 | 0.162 | 0.188 | 57% |
| SM d18:0/18:0 | 7,032.74 | 6,030.26 | 10,154.18 | 13,865.24 | 0.191 | 0.194 | 44% |
| SM d18:1/19:1 | 684.52 | 845.61 | 1,210.39 | 2,119.81 | 0.127 | 0.139 | 77% |
| SM d18:1/19:0 | 2,545.16 | 3,091.95 | 3,878.42 | 6,239.70 | 0.224 | 0.275 | 52% |
| SM d18:1/20:1 | 12,017.58 | 13,461.09 | 20,731.21 | 35,472.11 | 0.128 | 0.242 | 73% |
| SM d18:1/20:0 | 34,826.78 | 38,237.52 | 52,655.08 | 90,131.98 | 0.248 | 0.562 | 51% |
| SM d18:1/21:1 | 1,429.28 | 1,537.62 | 2,423.55 | 3,745.42 | 0.104 | 0.052 | 70% |
| SM d18:1/21:0 | 7,361.64 | 8,314.51 | 10,675.81 | 18,391.14 | 0.297 | 0.653 | 45% |
| SM d18:1/22:1 | 46,827.61 | 49,136.37 | 73,389.35 | 127,557.32 | 0.219 | 0.494 | 57% |
| SM d18:1/22:0 | 74,840.63 | 73,026.20 | 106,707.91 | 179,906.53 | 0.298 | 0.856 | 43% |
| SM d18:1/23:1 | 18,081.48 | 21,213.37 | 28,329.23 | 47,768.35 | 0.214 | 0.287 | 57% |
| SM d18:1/23:0 | 26,908.03 | 29,048.26 | 38,383.33 | 60,875.30 | 0.279 | 0.527 | 43% |
| SM d18:1/24:1 | 157,301.00 | 171,533.25 | 212,925.04 | 327,984.16 | 0.338 | 0.609 | 35% |
| SM d18:1/24:0 | 65,451.44 | 61,345.47 | 86,713.95 | 141,494.70 | 0.381 | 0.802 | 32% |
| SM d18:0/24:0 | 6,869.94 | 5,980.68 | 8,853.64 | 14,267.69 | 0.415 | 0.634 | 29% |
| SM d18:1/25:1 | 3,632.10 | 4,145.74 | 4,844.12 | 6,596.36 | 0.319 | 0.467 | 33% |
| SM d18:1/25:0 | 2,952.53 | 3,086.90 | 4,099.75 | 6,241.02 | 0.294 | 0.762 | 39% |
| SM d18:0/25:0 | 1,719.97 | 1,898.90 | 2,537.14 | 4,147.75 | 0.255 | 0.579 | 48% |
| SM d18:1/26:1 | 1,076.02 | 1,129.34 | 1,386.10 | 1,942.26 | 0.377 | 0.808 | 29% |
| SM d18:1/26:0 | 548.73 | 486.92 | 702.80 | 921.36 | 0.346 | 0.835 | 28% |
| SM d18:0/26:0 | 236.94 | 219.39 | 316.85 | 422.50 | 0.266 | 0.849 | 34% |
| Cer d18:1/16:0 | 733.94 | 326.00 | 723.88 | 286.79 | 0.879 | 0.924 | -1% |
| Cer d18:1/18:0 | 424.96 | 373.33 | 362.53 | 216.39 | 0.338 | 0.467 | -15% |
| Cer d18:0/18:0 | 252.42 | 239.77 | 250.46 | 198.62 | 0.967 | 0.457 | -1% |
| Cer d18:1/20:0 | 359.80 | 392.06 | 278.40 | 247.02 | 0.246 | 0.1 | -23% |
| Cer d18:1/22:0 | 997.55 | 430.14 | 1,056.45 | 515.93 | 0.57 | 0.703 | 6% |
| Cer d18:1/24:1 | 2,181.16 | 1,118.76 | 2,248.25 | 1,577.70 | 0.823 | 0.835 | 3% |
| Cer d18:1/24:0 | 3,080.36 | 1,228.06 | 3,046.26 | 1,635.38 | 0.914 | 0.489 | -1% |
| Cer d18:0/24:1 | 209.60 | 178.50 | 214.47 | 171.01 | 0.898 | 0.924 | 2% |
| GluCer d18:1/16:0 | 750.59 | 316.45 | 761.92 | 237.13 | 0.85 | 0.603 | 2% |
| GluCer d18:0/16:0 | 27.41 | 9.86 | 26.78 | 8.36 | 0.747 | 0.788 | -2% |
| GluCer d18:1/18:0 | 94.80 | 36.00 | 110.10 | 41.65 | 0.074 | 0.089 | 16% |
| GluCer d18:0/18:0 | 6.98 | 4.18 | 8.03 | 2.97 | 0.181 | **0.039** | 15% |
| GluCer d18:1/20:0 | 84.58 | 33.41 | 92.00 | 29.93 | 0.281 | 0.315 | 9% |
| GluCer d18:0/20:0 | 6.69 | 3.12 | 6.28 | 3.67 | 0.586 | 0.268 | -6% |
| GluCer d18:1/22:0 | 732.92 | 310.43 | 725.66 | 274.95 | 0.909 | 0.931 | -1% |
| GluCer d18:0/22:0 | 17.73 | 7.74 | 18.76 | 7.80 | 0.541 | 0.71 | 6% |
| GluCer d18:1/24:1 | 930.28 | 317.27 | 984.06 | 283.66 | 0.409 | 0.446 | 6% |
| GluCer d18:0/24:1 | 963.37 | 420.63 | 914.51 | 371.85 | 0.569 | 0.516 | -5% |
| GluCer d18:1/24:0 | 26.80 | 12.40 | 27.27 | 9.93 | 0.846 | 0.562 | 2% |
| GluCer d18:0/24:0 | 22.61 | 10.17 | 21.22 | 8.83 | 0.502 | 0.373 | -6% |
| LacCer d18:1/16:0 | 3,319.71 | 1,121.12 | 3,200.51 | 834.62 | 0.582 | 0.965 | -4% |
| LacCer d18:0/16:0 | 78.57 | 24.46 | 76.37 | 21.47 | 0.658 | 0.749 | -3% |
| LacCer d18:1/18:0 | 100.88 | 33.04 | 101.01 | 63.47 | 0.991 | 0.377 | 0% |
| LacCer d18:1/20:0 | 40.34 | 17.09 | 42.25 | 24.07 | 0.677 | 0.924 | 5% |
| LacCer d18:1/22:0 | 181.20 | 73.18 | 176.46 | 97.89 | 0.802 | 0.436 | -3% |
| LacCer d18:1/24:1 | 660.40 | 251.70 | 638.00 | 222.82 | 0.663 | 0.775 | -3% |
| LacCer d18:1/24:0 | 266.52 | 105.28 | 267.49 | 136.44 | 0.971 | 0.938 | 0% |
| LacCer d18:0/24:1 | 18.75 | 8.79 | 16.06 | 6.37 | 0.104 | 0.203 | -14% |
| Gb3 d18:1/16:0 | 1,412.52 | 378.84 | 1,452.58 | 326.65 | 0.6 | 0.483 | 3% |
| Gb3 d18:1/18:0 | 160.89 | 51.28 | 167.72 | 56.60 | 0.561 | 0.755 | 4% |
| Gb3 d18:1/20:0 | 63.95 | 26.70 | 74.61 | 23.52 | 0.052 | **0.009** | 17% |
| Gb3 d18:1/22:0 | 206.38 | 70.35 | 224.67 | 78.68 | 0.262 | 0.332 | 9% |
| Gb3 d18:1/24:1 | 27.45 | 15.06 | 25.83 | 14.33 | 0.611 | 0.671 | -6% |
| Gb3 d18:1/24:0 | 81.59 | 28.43 | 77.97 | 31.65 | 0.58 | 0.467 | -4% |
| GM3 d18:1/16:0 | 3,403.98 | 967.45 | 3,295.38 | 790.38 | 0.568 | 0.788 | -3% |
| GM3 d18:0/16:0 | 723.50 | 220.45 | 686.62 | 163.95 | 0.377 | 0.849 | -5% |
| GM3 d18:1/18:1 | 387.44 | 119.25 | 359.98 | 123.08 | 0.298 | 0.246 | -7% |
| GM3 d18:1/18:0 | 1,559.14 | 427.58 | 1,301.05 | 362.73 | **0.003** | **0.003** | -17% |
| GM3 d18:0/18:0 | 562.20 | 125.24 | 508.78 | 141.29 | 0.069 | **0.038** | -10% |
| GM3 d18:1/20:1 | 194.20 | 67.06 | 188.98 | 67.63 | 0.721 | 0.723 | -3% |
| GM3 d18:1/20:0 | 698.33 | 251.91 | 594.50 | 191.91 | **0.037** | **0.045** | -15% |
| GM3 d18:0/20:0 | 191.02 | 71.34 | 168.85 | 69.51 | 0.149 | 0.109 | -12% |
| GM3 d18:1/22:1 | 873.83 | 285.60 | 828.96 | 233.36 | 0.425 | 0.808 | -5% |
| GM3 d18:1/22:0 | 1,949.95 | 689.83 | 1,735.82 | 565.09 | 0.117 | 0.13 | -11% |
| GM3 d18:0/22:0 | 531.34 | 214.52 | 465.81 | 211.41 | 0.158 | 0.096 | -12% |
| GM3 d18:1/24:1 | 2,522.35 | 821.36 | 2,343.51 | 729.56 | 0.288 | 0.337 | -7% |
| GM3 d18:1/24:0 | 1,938.94 | 625.09 | 1,683.28 | 456.44 | **0.036** | 0.06 | -13% |
| GM3 d18:0/24:0 | 477.69 | 137.55 | 466.07 | 138.48 | 0.698 | 0.622 | -2% |
| FFA | 10,700,169.57 | 895,473.03 | 10,623,454.39 | 947,850.78 | 0.702 | 0.634 | -1% |
| SM | 870,614.80 | 928,565.89 | 1,314,656.93 | 2,215,685.17 | 0.241 | 0.544 | 51% |
| Cer | 8,239.79 | 2,722.51 | 8,180.70 | 3,918.70 | 0.936 | 0.426 | -1% |
| GluCer | 3,664.76 | 1,394.08 | 3,696.58 | 1,189.34 | 0.909 | 0.869 | 1% |
| LacCer | 4,666.37 | 1,536.55 | 4,518.14 | 1,260.03 | 0.624 | 0.89 | -3% |
| GM3 | 16,013.92 | 4,316.51 | 14,627.60 | 3,612.06 | 0.109 | 0.136 | -9% |
| Gb3 | 1,952.78 | 499.64 | 2,023.37 | 471.45 | 0.502 | 0.653 | 4% |
| SM/Cer | 108.53 | 114.39 | 180.94 | 301.07 | 0.136 | 0.539 | 67% |

**Supplementary Table 2.** Age differences in individual lipid species

| Lipids | Age ≤ 26 (n=43) |  | Age > 26 (n=43) |  | P value |  | Percentage changed in age |
| --- | --- | --- | --- | --- | --- | --- | --- |
|  | Average lipid concentration (nM) | Standard deviation | Average lipid concentration (nM) | Standard deviation | t.test | Man whitney u test |  |
| FFA22:6 | 25,664.85 | 10,909.68 | 30,151.73 | 10,450.19 | 0.055 | **0.009** | 17% |
| FFA22:5 | 32,676.18 | 4,628.27 | 37,258.53 | 5,829.72 | **0.000** | **0.000** | 14% |
| FFA22:4 | 44,495.41 | 3,982.61 | 47,188.29 | 5,344.60 | **0.010** | **0.023** | 6% |
| FFA20:5 | 37,143.60 | 7,624.72 | 38,654.07 | 5,725.95 | 0.302 | **0.049** | 4% |
| FFA20:4 | 69,792.26 | 29,525.78 | 73,715.15 | 31,512.28 | 0.553 | 0.113 | 6% |
| FFA20:3 | 70,432.19 | 5,427.57 | 73,817.44 | 6,730.52 | **0.012** | **0.008** | 5% |
| FFA18:3 | 19,424.87 | 2,551.20 | 21,974.15 | 4,353.13 | **0.001** | **0.001** | 13% |
| FFA18:2 | 694,918.69 | 193,721.87 | 890,918.47 | 271,398.54 | **0.000** | **0.000** | 28% |
| FFA18:1 | 330,532.70 | 53,962.63 | 390,608.76 | 75,812.34 | **0.000** | **0.000** | 18% |
| FFA18:0 | 2,406,859.07 | 237,006.03 | 2,533,686.06 | 199,862.86 | **0.009** | **0.010** | 5% |
| FFA16:1 | 838,192.23 | 270,514.95 | 926,880.34 | 270,375.93 | 0.132 | 0.111 | 11% |
| FFA16:0 | 2,596,906.98 | 213,413.67 | 2,716,354.58 | 195,905.92 | **0.008** | **0.004** | 5% |
| FFA14:0 | 225,165.36 | 20,514.97 | 234,591.00 | 20,035.65 | **0.034** | 0.073 | 4% |
| FA13:0 | 1,383,897.97 | 184,211.09 | 1,390,041.25 | 169,360.87 | 0.872 | 0.65 | 0% |
| FA12:0 | 1,574,468.00 | 218,118.19 | 1,561,861.55 | 205,569.19 | 0.783 | 0.969 | -1% |
| SM d18:1/16:1 | 47,803.64 | 70,685.87 | 30,754.08 | 24,393.51 | 0.139 | 0.323 | -36% |
| SM d18:1/16:0 | 448,597.44 | 798,519.85 | 248,945.51 | 290,577.87 | 0.127 | 0.2 | -45% |
| SM d18:0/16:0 | 46,260.37 | 83,806.11 | 25,994.88 | 28,998.68 | 0.138 | 0.222 | -44% |
| SM d18:1/17:1 | 1,207.23 | 2,033.47 | 728.37 | 888.38 | 0.161 | 0.263 | -40% |
| SM d18:1/17:0 | 8,352.34 | 14,503.31 | 4,846.46 | 6,359.45 | 0.150 | 0.171 | -42% |
| SM d18:1/18:1 | 39,237.53 | 64,227.32 | 23,825.31 | 24,395.07 | 0.145 | 0.185 | -39% |
| SM d18:1/18:0 | 81,892.35 | 131,904.76 | 50,689.99 | 49,775.32 | 0.150 | 0.245 | -38% |
| SM d18:0/18:0 | 10,140.52 | 14,339.18 | 7,264.17 | 5,904.19 | 0.227 | 0.694 | -28% |
| SM d18:1/19:1 | 1,255.75 | 2,181.41 | 675.86 | 829.90 | 0.107 | 0.061 | -46% |
| SM d18:1/19:0 | 3,973.47 | 6,438.94 | 2,543.12 | 2,998.39 | 0.190 | 0.336 | -36% |
| SM d18:1/20:1 | 21,452.24 | 36,504.16 | 11,904.48 | 13,261.38 | 0.111 | 0.121 | -45% |
| SM d18:1/20:0 | 54,530.10 | 92,935.16 | 34,195.59 | 37,081.26 | 0.186 | 0.385 | -37% |
| SM d18:1/21:1 | 2,442.31 | 3,865.52 | 1,479.89 | 1,520.77 | 0.132 | 0.188 | -39% |
| SM d18:1/21:0 | 11,098.66 | 18,995.34 | 7,170.01 | 7,966.07 | 0.215 | 0.487 | -35% |
| SM d18:1/22:1 | 77,117.32 | 131,391.34 | 44,952.78 | 47,340.52 | 0.135 | 0.132 | -42% |
| SM d18:1/22:0 | 113,955.80 | 185,271.33 | 69,816.03 | 69,459.72 | 0.147 | 0.168 | -39% |
| SM d18:1/23:1 | 29,840.28 | 49,233.44 | 17,285.38 | 20,286.50 | 0.126 | 0.141 | -42% |
| SM d18:1/23:0 | 40,411.11 | 62,807.51 | 25,680.85 | 27,663.27 | 0.163 | 0.218 | -36% |
| SM d18:1/24:1 | 225,164.02 | 337,144.44 | 148,942.76 | 165,876.97 | 0.187 | 0.121 | -34% |
| SM d18:1/24:0 | 93,791.48 | 145,581.58 | 59,857.34 | 58,241.73 | 0.160 | 0.197 | -36% |
| SM d18:0/24:0 | 9,596.96 | 14,682.48 | 6,265.02 | 5,662.81 | 0.169 | 0.302 | -35% |
| SM d18:1/25:1 | 5,074.90 | 6,751.28 | 3,485.87 | 4,056.53 | 0.189 | **0.046** | -31% |
| SM d18:1/25:0 | 4,351.82 | 6,397.48 | 2,780.50 | 3,009.93 | 0.149 | 0.086 | -36% |
| SM d18:0/25:0 | 2,703.21 | 4,256.07 | 1,610.91 | 1,842.91 | 0.126 | 0.065 | -40% |
| SM d18:1/26:1 | 1,476.26 | 1,987.36 | 1,007.49 | 1,101.38 | 0.180 | 0.07 | -32% |
| SM d18:1/26:0 | 774.38 | 949.39 | 487.90 | 445.45 | 0.077 | 0.123 | -37% |
| SM d18:0/26:0 | 347.91 | 429.81 | 211.46 | 211.98 | 0.065 | 0.07 | -39% |
| Cer d18:1/16:0 | 733.36 | 278.67 | 723.76 | 330.39 | 0.885 | 0.434 | -1% |
| Cer d18:1/18:0 | 396.83 | 217.33 | 386.31 | 366.45 | 0.872 | 0.228 | -3% |
| Cer d18:0/18:0 | 250.92 | 195.82 | 251.82 | 239.39 | 0.985 | 0.543 | 0% |
| Cer d18:1/20:0 | 288.39 | 180.52 | 344.14 | 421.08 | 0.427 | 0.839 | 19% |
| Cer d18:1/22:0 | 994.94 | 515.38 | 1,063.16 | 436.77 | 0.510 | 0.235 | 7% |
| Cer d18:1/24:1 | 1,884.11 | 859.78 | 2,549.98 | 1,692.71 | **0.024** | **0.038** | 35% |
| Cer d18:1/24:0 | 2,705.28 | 1,173.58 | 3,418.96 | 1,621.26 | **0.022** | **0.012** | 26% |
| Cer d18:0/24:1 | 199.25 | 128.68 | 225.16 | 209.82 | 0.492 | 0.4 | 13% |
| GluCer d18:1/16:0 | 780.49 | 293.25 | 732.81 | 257.19 | 0.425 | 0.694 | -6% |
| GluCer d18:0/16:0 | 27.49 | 9.59 | 26.66 | 8.55 | 0.674 | 0.806 | -3% |
| GluCer d18:1/18:0 | 102.43 | 40.86 | 103.53 | 38.87 | 0.898 | 0.575 | 1% |
| GluCer d18:0/18:0 | 8.07 | 3.61 | 7.01 | 3.55 | 0.172 | 0.245 | -13% |
| GluCer d18:1/20:0 | 88.98 | 34.66 | 88.12 | 28.69 | 0.900 | 0.894 | -1% |
| GluCer d18:0/20:0 | 6.53 | 3.36 | 6.41 | 3.50 | 0.874 | 0.792 | -2% |
| GluCer d18:1/22:0 | 747.40 | 327.65 | 710.68 | 249.93 | 0.561 | 0.931 | -5% |
| GluCer d18:0/22:0 | 19.35 | 8.13 | 17.21 | 7.27 | 0.202 | 0.308 | -11% |
| GluCer d18:1/24:1 | 982.35 | 324.90 | 935.74 | 272.91 | 0.473 | 0.839 | -5% |
| GluCer d18:0/24:1 | 961.44 | 444.42 | 913.03 | 339.03 | 0.572 | 0.99 | -5% |
| GluCer d18:1/24:0 | 27.18 | 11.99 | 26.92 | 10.22 | 0.912 | 0.863 | -1% |
| GluCer d18:0/24:0 | 23.29 | 10.37 | 20.44 | 8.30 | 0.162 | 0.284 | -12% |
| LacCer d18:1/16:0 | 3,116.73 | 927.56 | 3,395.17 | 1,010.28 | 0.187 | 0.158 | 9% |
| LacCer d18:0/16:0 | 75.82 | 24.32 | 78.97 | 21.33 | 0.524 | 0.429 | 4% |
| LacCer d18:1/18:0 | 104.41 | 64.03 | 97.49 | 34.72 | 0.535 | 0.736 | -7% |
| LacCer d18:1/20:0 | 42.88 | 25.26 | 39.85 | 15.85 | 0.508 | 0.789 | -7% |
| LacCer d18:1/22:0 | 178.95 | 103.59 | 178.38 | 67.23 | 0.976 | 0.492 | 0% |
| LacCer d18:1/24:1 | 660.78 | 255.14 | 636.05 | 216.49 | 0.629 | 0.894 | -4% |
| LacCer d18:1/24:0 | 271.48 | 146.86 | 262.59 | 92.89 | 0.738 | 0.812 | -3% |
| LacCer d18:0/24:1 | 17.04 | 6.83 | 17.58 | 8.49 | 0.744 | 0.832 | 3% |
| Gb3 d18:1/16:0 | 1,398.17 | 314.51 | 1,469.74 | 383.23 | 0.347 | 0.218 | 5% |
| Gb3 d18:1/18:0 | 160.73 | 53.32 | 168.35 | 55.00 | 0.516 | 0.442 | 5% |
| Gb3 d18:1/20:0 | 71.47 | 24.72 | 67.83 | 26.36 | 0.510 | 0.344 | -5% |
| Gb3 d18:1/22:0 | 213.78 | 69.17 | 218.54 | 81.25 | 0.771 | 0.969 | 2% |
| Gb3 d18:1/24:1 | 26.62 | 10.46 | 26.54 | 17.96 | 0.980 | 0.252 | 0% |
| Gb3 d18:1/24:0 | 80.69 | 29.30 | 78.62 | 31.14 | 0.752 | 0.887 | -3% |
| GM3 d18:1/16:0 | 3,164.27 | 745.19 | 3,527.52 | 959.98 | 0.053 | 0.059 | 11% |
| GM3 d18:0/16:0 | 656.82 | 172.45 | 750.74 | 200.90 | **0.022** | **0.027** | 14% |
| GM3 d18:1/18:1 | 352.47 | 114.04 | 393.04 | 126.39 | 0.122 | 0.121 | 12% |
| GM3 d18:1/18:0 | 1,331.05 | 361.03 | 1,511.14 | 444.67 | **0.042** | 0.088 | 14% |
| GM3 d18:0/18:0 | 518.18 | 123.86 | 549.07 | 146.89 | 0.295 | 0.455 | 6% |
| GM3 d18:1/20:1 | 182.87 | 73.94 | 199.95 | 58.94 | 0.240 | **0.047** | 9% |
| GM3 d18:1/20:0 | 582.33 | 206.64 | 703.26 | 231.72 | **0.012** | **0.009** | 21% |
| GM3 d18:0/20:0 | 168.35 | 69.42 | 189.97 | 71.38 | 0.158 | 0.102 | 13% |
| GM3 d18:1/22:1 | 785.34 | 231.35 | 914.33 | 270.32 | **0.020** | **0.026** | 16% |
| GM3 d18:1/22:0 | 1,669.14 | 552.09 | 2,001.69 | 667.80 | **0.014** | **0.011** | 20% |
| GM3 d18:0/22:0 | 430.96 | 182.68 | 561.61 | 225.16 | **0.004** | **0.001** | 30% |
| GM3 d18:1/24:1 | 2,292.58 | 758.54 | 2,560.80 | 775.11 | 0.109 | 0.098 | 12% |
| GM3 d18:1/24:0 | 1,697.03 | 521.51 | 1,907.36 | 569.85 | 0.078 | 0.059 | 12% |
| GM3 d18:0/24:0 | 435.18 | 133.48 | 507.77 | 132.93 | **0.013** | **0.010** | 17% |
| FFA | 10,350,570.37 | 866,866.65 | 10,967,701.37 | 873,939.57 | **0.001** | **0.003** | 6% |
| SM | 1,382,849.40 | 2,282,469.10 | 833,402.02 | 892,639.92 | 0.145 | 0.191 | -40% |
| Cer | 7,453.08 | 2,426.09 | 8,963.28 | 4,035.35 | **0.038** | **0.035** | 20% |
| GluCer | 3,775.01 | 1,434.61 | 3,588.55 | 1,115.71 | 0.503 | 0.983 | -5% |
| LacCer | 4,468.08 | 1,426.28 | 4,706.09 | 1,356.97 | 0.430 | 0.271 | 5% |
| GM3 | 14,266.55 | 3,661.59 | 16,278.25 | 4,096.39 | **0.019** | **0.019** | 14% |
| Gb3 | 1,951.46 | 441.57 | 2,029.62 | 523.81 | 0.456 | 0.323 | 4% |
| SM/Cer | 192.98 | 307.62 | 101.54 | 114.15 | 0.071 | **0.005** | -47% |

**Supplementary Table 3.**
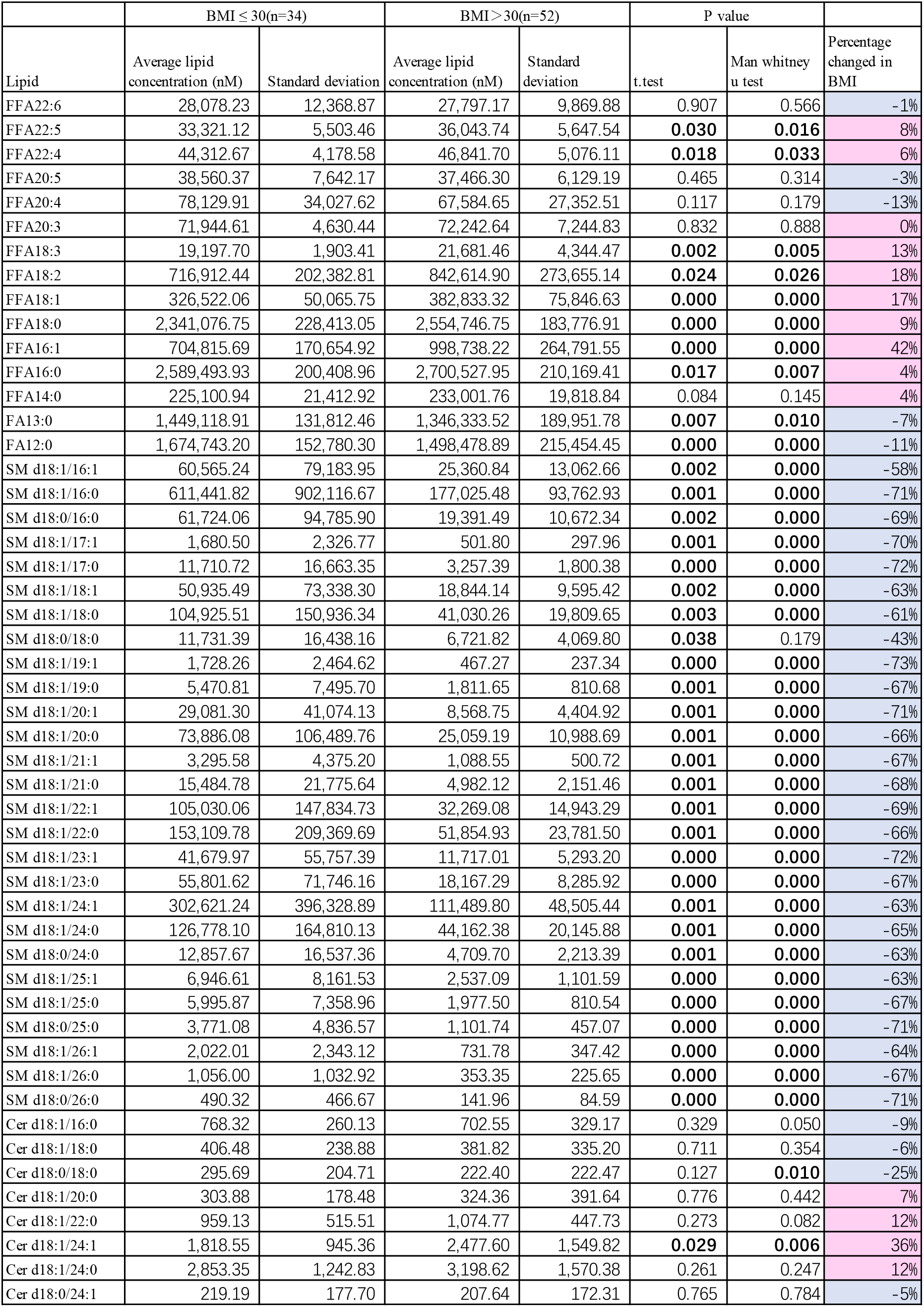
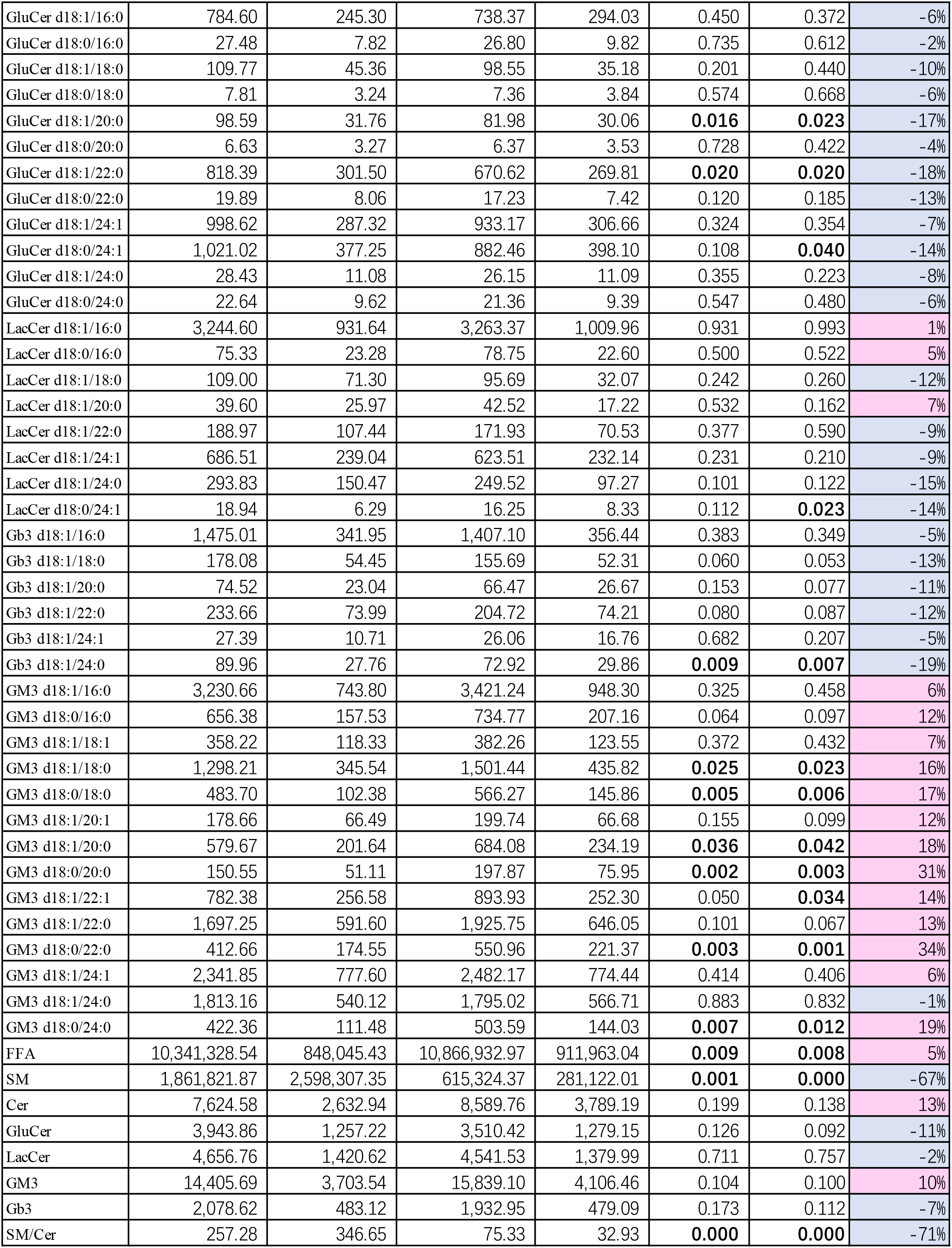
BMI differences in individual lipid species

| Lipid | BMI ≤ 30(n=34) |  | BMI > 30(n=52) |  | P value |  | Percentage changed in BMI |
| --- | --- | --- | --- | --- | --- | --- | --- |
|  | Average lipid concentration (nM) | Standard deviation | Average lipid concentration (nM) | Standard deviation | t.test | Man whitney u test |  |
| FFA22:6 | 28,078.23 | 12,368.87 | 27,797.17 | 9,869.88 | 0.907 | 0.566 | -1% |
| FFA22:5 | 33,321.12 | 5,503.46 | 36,043.74 | 5,647.54 | **0.030** | **0.016** | 8% |
| FFA22:4 | 44,312.67 | 4,178.58 | 46,841.70 | 5,076.11 | **0.018** | **0.033** | 6% |
| FFA20:5 | 38,560.37 | 7,642.17 | 37,466.30 | 6,129.19 | 0.465 | 0.314 | -3% |
| FFA20:4 | 78,129.91 | 34,027.62 | 67,584.65 | 27,352.51 | 0.117 | 0.179 | -13% |
| FFA20:3 | 71,944.61 | 4,630.44 | 72,242.64 | 7,244.83 | 0.832 | 0.888 | 0% |
| FFA18:3 | 19,197.70 | 1,903.41 | 21,681.46 | 4,344.47 | **0.002** | **0.005** | 13% |
| FFA18:2 | 716,912.44 | 202,382.81 | 842,614.90 | 273,655.14 | **0.024** | **0.026** | 18% |
| FFA18:1 | 326,522.06 | 50,065.75 | 382,833.32 | 75,846.63 | **0.000** | **0.000** | 17% |
| FFA18:0 | 2,341,076.75 | 228,413.05 | 2,554,746.75 | 183,776.91 | **0.000** | **0.000** | 9% |
| FFA16:1 | 704,815.69 | 170,654.92 | 998,738.22 | 264,791.55 | **0.000** | **0.000** | 42% |
| FFA16:0 | 2,589,493.93 | 200,408.96 | 2,700,527.95 | 210,169.41 | **0.017** | **0.007** | 4% |
| FFA14:0 | 225,100.94 | 21,412.92 | 233,001.76 | 19,818.84 | 0.084 | 0.145 | 4% |
| FA13:0 | 1,449,118.91 | 131,812.46 | 1,346,333.52 | 189,951.78 | **0.007** | **0.010** | -7% |
| FA12:0 | 1,674,743.20 | 152,780.30 | 1,498,478.89 | 215,454.45 | **0.000** | **0.000** | -11% |
| SM d18:1/16:1 | 60,565.24 | 79,183.95 | 25,360.84 | 13,062.66 | **0.002** | **0.000** | -58% |
| SM d18:1/16:0 | 611,441.82 | 902,116.67 | 177,025.48 | 93,762.93 | **0.001** | **0.000** | -71% |
| SM d18:0/16:0 | 61,724.06 | 94,785.90 | 19,391.49 | 10,672.34 | **0.002** | **0.000** | -69% |
| SM d18:1/17:1 | 1,680.50 | 2,326.77 | 501.80 | 297.96 | **0.001** | **0.000** | -70% |
| SM d18:1/17:0 | 11,710.72 | 16,663.35 | 3,257.39 | 1,800.38 | **0.000** | **0.000** | -72% |
| SM d18:1/18:1 | 50,935.49 | 73,338.30 | 18,844.14 | 9,595.42 | **0.002** | **0.000** | -63% |
| SM d18:1/18:0 | 104,925.51 | 150,936.34 | 41,030.26 | 19,809.65 | **0.003** | **0.000** | -61% |
| SM d18:0/18:0 | 11,731.39 | 16,438.16 | 6,721.82 | 4,069.80 | **0.038** | 0.179 | -43% |
| SM d18:1/19:1 | 1,728.26 | 2,464.62 | 467.27 | 237.34 | **0.000** | **0.000** | -73% |
| SM d18:1/19:0 | 5,470.81 | 7,495.70 | 1,811.65 | 810.68 | **0.001** | **0.000** | -67% |
| SM d18:1/20:1 | 29,081.30 | 41,074.13 | 8,568.75 | 4,404.92 | **0.001** | **0.000** | -71% |
| SM d18:1/20:0 | 73,886.08 | 106,489.76 | 25,059.19 | 10,988.69 | **0.001** | **0.000** | -66% |
| SM d18:1/21:1 | 3,295.58 | 4,375.20 | 1,088.55 | 500.72 | **0.001** | **0.000** | -67% |
| SM d18:1/21:0 | 15,484.78 | 21,775.64 | 4,982.12 | 2,151.46 | **0.001** | **0.000** | -68% |
| SM d18:1/22:1 | 105,030.06 | 147,834.73 | 32,269.08 | 14,943.29 | **0.001** | **0.000** | -69% |
| SM d18:1/22:0 | 153,109.78 | 209,369.69 | 51,854.93 | 23,781.50 | **0.001** | **0.000** | -66% |
| SM d18:1/23:1 | 41,679.97 | 55,757.39 | 11,717.01 | 5,293.20 | **0.000** | **0.000** | -72% |
| SM d18:1/23:0 | 55,801.62 | 71,746.16 | 18,167.29 | 8,285.92 | **0.000** | **0.000** | -67% |
| SM d18:1/24:1 | 302,621.24 | 396,328.89 | 111,489.80 | 48,505.44 | **0.001** | **0.000** | -63% |
| SM d18:1/24:0 | 126,778.10 | 164,810.13 | 44,162.38 | 20,145.88 | **0.001** | **0.000** | -65% |
| SM d18:0/24:0 | 12,857.67 | 16,537.36 | 4,709.70 | 2,213.39 | **0.001** | **0.000** | -63% |
| SM d18:1/25:1 | 6,946.61 | 8,161.53 | 2,537.09 | 1,101.59 | **0.000** | **0.000** | -63% |
| SM d18:1/25:0 | 5,995.87 | 7,358.96 | 1,977.50 | 810.54 | **0.000** | **0.000** | -67% |
| SM d18:0/25:0 | 3,771.08 | 4,836.57 | 1,101.74 | 457.07 | **0.000** | **0.000** | -71% |
| SM d18:1/26:1 | 2,022.01 | 2,343.12 | 731.78 | 347.42 | **0.000** | **0.000** | -64% |
| SM d18:1/26:0 | 1,056.00 | 1,032.92 | 353.35 | 225.65 | **0.000** | **0.000** | -67% |
| SM d18:0/26:0 | 490.32 | 466.67 | 141.96 | 84.59 | **0.000** | **0.000** | -71% |
| Cer d18:1/16:0 | 768.32 | 260.13 | 702.55 | 329.17 | 0.329 | 0.050 | -9% |
| Cer d18:1/18:0 | 406.48 | 238.88 | 381.82 | 335.20 | 0.711 | 0.354 | -6% |
| Cer d18:0/18:0 | 295.69 | 204.71 | 222.40 | 222.47 | 0.127 | **0.010** | -25% |
| Cer d18:1/20:0 | 303.88 | 178.48 | 324.36 | 391.64 | 0.776 | 0.442 | 7% |
| Cer d18:1/22:0 | 959.13 | 515.51 | 1,074.77 | 447.73 | 0.273 | 0.082 | 12% |
| Cer d18:1/24:1 | 1,818.55 | 945.36 | 2,477.60 | 1,549.82 | **0.029** | **0.006** | 36% |
| Cer d18:1/24:0 | 2,853.35 | 1,242.83 | 3,198.62 | 1,570.38 | 0.261 | 0.247 | 12% |
| Cer d18:0/24:1 | 219.19 | 177.70 | 207.64 | 172.31 | 0.765 | 0.784 | -5% |
| GluCer d18:1/16:0 | 784.60 | 245.30 | 738.37 | 294.03 | 0.450 | 0.372 | -6% |
| GluCer d18:0/16:0 | 27.48 | 7.82 | 26.80 | 9.82 | 0.735 | 0.612 | -2% |
| GluCer d18:1/18:0 | 109.77 | 45.36 | 98.55 | 35.18 | 0.201 | 0.440 | -10% |
| GluCer d18:0/18:0 | 7.81 | 3.24 | 7.36 | 3.84 | 0.574 | 0.668 | -6% |
| GluCer d18:1/20:0 | 98.59 | 31.76 | 81.98 | 30.06 | **0.016** | **0.023** | -17% |
| GluCer d18:0/20:0 | 6.63 | 3.27 | 6.37 | 3.53 | 0.728 | 0.422 | -4% |
| GluCer d18:1/22:0 | 818.39 | 301.50 | 670.62 | 269.81 | **0.020** | **0.020** | -18% |
| GluCer d18:0/22:0 | 19.89 | 8.06 | 17.23 | 7.42 | 0.120 | 0.185 | -13% |
| GluCer d18:1/24:1 | 998.62 | 287.32 | 933.17 | 306.66 | 0.324 | 0.354 | -7% |
| GluCer d18:0/24:1 | 1,021.02 | 377.25 | 882.46 | 398.10 | 0.108 | **0.040** | -14% |
| GluCer d18:1/24:0 | 28.43 | 11.08 | 26.15 | 11.09 | 0.355 | 0.223 | -8% |
| GluCer d18:0/24:0 | 22.64 | 9.62 | 21.36 | 9.39 | 0.547 | 0.480 | -6% |
| LacCer d18:1/16:0 | 3,244.60 | 931.64 | 3,263.37 | 1,009.96 | 0.931 | 0.993 | 1% |
| LacCer d18:0/16:0 | 75.33 | 23.28 | 78.75 | 22.60 | 0.500 | 0.522 | 5% |
| LacCer d18:1/18:0 | 109.00 | 71.30 | 95.69 | 32.07 | 0.242 | 0.260 | -12% |
| LacCer d18:1/20:0 | 39.60 | 25.97 | 42.52 | 17.22 | 0.532 | 0.162 | 7% |
| LacCer d18:1/22:0 | 188.97 | 107.44 | 171.93 | 70.53 | 0.377 | 0.590 | -9% |
| LacCer d18:1/24:1 | 686.51 | 239.04 | 623.51 | 232.14 | 0.231 | 0.210 | -9% |
| LacCer d18:1/24:0 | 293.83 | 150.47 | 249.52 | 97.27 | 0.101 | 0.122 | -15% |
| LacCer d18:0/24:1 | 18.94 | 6.29 | 16.25 | 8.33 | 0.112 | **0.023** | -14% |
| Gb3 d18:1/16:0 | 1,475.01 | 341.95 | 1,407.10 | 356.44 | 0.383 | 0.349 | -5% |
| Gb3 d18:1/18:0 | 178.08 | 54.45 | 155.69 | 52.31 | 0.060 | 0.053 | -13% |
| Gb3 d18:1/20:0 | 74.52 | 23.04 | 66.47 | 26.67 | 0.153 | 0.077 | -11% |
| Gb3 d18:1/22:0 | 233.66 | 73.99 | 204.72 | 74.21 | 0.080 | 0.087 | -12% |
| Gb3 d18:1/24:1 | 27.39 | 10.71 | 26.06 | 16.76 | 0.682 | 0.207 | -5% |
| Gb3 d18:1/24:0 | 89.96 | 27.76 | 72.92 | 29.86 | **0.009** | **0.007** | -19% |
| GM3 d18:1/16:0 | 3,230.66 | 743.80 | 3,421.24 | 948.30 | 0.325 | 0.458 | 6% |
| GM3 d18:0/16:0 | 656.38 | 157.53 | 734.77 | 207.16 | 0.064 | 0.097 | 12% |
| GM3 d18:1/18:1 | 358.22 | 118.33 | 382.26 | 123.55 | 0.372 | 0.432 | 7% |
| GM3 d18:1/18:0 | 1,298.21 | 345.54 | 1,501.44 | 435.82 | **0.025** | **0.023** | 16% |
| GM3 d18:0/18:0 | 483.70 | 102.38 | 566.27 | 145.86 | **0.005** | **0.006** | 17% |
| GM3 d18:1/20:1 | 178.66 | 66.49 | 199.74 | 66.68 | 0.155 | 0.099 | 12% |
| GM3 d18:1/20:0 | 579.67 | 201.64 | 684.08 | 234.19 | **0.036** | **0.042** | 18% |
| GM3 d18:0/20:0 | 150.55 | 51.11 | 197.87 | 75.95 | **0.002** | **0.003** | 31% |
| GM3 d18:1/22:1 | 782.38 | 256.58 | 893.93 | 252.30 | 0.050 | **0.034** | 14% |
| GM3 d18:1/22:0 | 1,697.25 | 591.60 | 1,925.75 | 646.05 | 0.101 | 0.067 | 13% |
| GM3 d18:0/22:0 | 412.66 | 174.55 | 550.96 | 221.37 | **0.003** | **0.001** | 34% |
| GM3 d18:1/24:1 | 2,341.85 | 777.60 | 2,482.17 | 774.44 | 0.414 | 0.406 | 6% |
| GM3 d18:1/24:0 | 1,813.16 | 540.12 | 1,795.02 | 566.71 | 0.883 | 0.832 | -1% |
| GM3 d18:0/24:0 | 422.36 | 111.48 | 503.59 | 144.03 | **0.007** | **0.012** | 19% |
| FFA | 10,341,328.54 | 848,045.43 | 10,866,932.97 | 911,963.04 | **0.009** | **0.008** | 5% |
| SM | 1,861,821.87 | 2,598,307.35 | 615,324.37 | 281,122.01 | **0.001** | **0.000** | -67% |
| Cer | 7,624.58 | 2,632.94 | 8,589.76 | 3,789.19 | 0.199 | 0.138 | 13% |
| GluCer | 3,943.86 | 1,257.22 | 3,510.42 | 1,279.15 | 0.126 | 0.092 | -11% |
| LacCer | 4,656.76 | 1,420.62 | 4,541.53 | 1,379.99 | 0.711 | 0.757 | -2% |
| GM3 | 14,405.69 | 3,703.54 | 15,839.10 | 4,106.46 | 0.104 | 0.100 | 10% |
| Gb3 | 2,078.62 | 483.12 | 1,932.95 | 479.09 | 0.173 | 0.112 | -7% |
| SM/Cer | 257.28 | 346.65 | 75.33 | 32.93 | **0.000** | **0.000** | -71% |

**Supplementary Table 4.** GIR differences in individual lipid species

| Lipid | GIR < 5.1 (n=42) |  | GIR ≥ 5.1 (n=44) |  | P value |  | Percentage changed in GIR |
| --- | --- | --- | --- | --- | --- | --- | --- |
|  | Average lipid concentration (nM) | Standard deviation | Average lipid concentration (nM) | Standard deviation | t.test | Man whitney u test |  |
| FFA22:6 | 26,607.05 | 9,332.87 | 29,150.38 | 12,112.08 | 0.280 | 0.622 | 10% |
| FFA22:5 | 35,920.88 | 5,979.61 | 34,057.17 | 5,365.49 | 0.132 | 0.154 | -5% |
| FFA22:4 | 46,854.61 | 5,184.60 | 44,875.12 | 4,408.02 | 0.059 | 0.104 | -4% |
| FFA20:5 | 37,059.56 | 5,921.87 | 38,699.97 | 7,427.44 | 0.262 | 0.142 | 4% |
| FFA20:4 | 65,459.54 | 25,719.95 | 77,761.77 | 33,507.37 | 0.060 | 0.104 | 19% |
| FFA20:3 | 72,590.15 | 7,563.17 | 71,680.63 | 4,878.05 | 0.507 | 0.762 | -1% |
| FFA18:3 | 21,879.83 | 4,642.97 | 19,572.84 | 2,213.94 | **0.004** | **0.023** | -11% |
| FFA18:2 | 825,883.82 | 287,515.05 | 761,451.75 | 216,593.93 | 0.242 | 0.407 | -8% |
| FFA18:1 | 382,384.30 | 78,245.13 | 339,748.69 | 59,301.88 | **0.005** | **0.007** | -11% |
| FFA18:0 | 2,545,189.69 | 172,275.66 | 2,398,760.76 | 250,820.40 | **0.002** | **0.004** | -6% |
| FFA16:1 | 1,000,727.24 | 245,056.04 | 769,717.65 | 250,768.36 | **0.000** | **0.000** | -23% |
| FFA16:0 | 2,689,712.49 | 204,045.72 | 2,625,052.78 | 217,529.74 | 0.159 | 0.106 | -2% |
| FFA14:0 | 232,674.32 | 19,646.13 | 227,209.14 | 21,555.62 | 0.223 | 0.325 | -2% |
| FA13:0 | 1,365,213.91 | 169,640.88 | 1,407,736.41 | 181,208.14 | 0.265 | 0.131 | 3% |
| FA12:0 | 1,519,191.44 | 201,698.34 | 1,614,912.05 | 210,845.36 | **0.034** | **0.016** | 6% |
| SM d18:1/16:1 | 25,301.54 | 13,156.13 | 52,620.85 | 71,195.78 | **0.017** | **0.002** | 108% |
| SM d18:1/16:0 | 175,643.25 | 91,966.50 | 514,030.24 | 812,355.35 | **0.009** | **0.000** | 193% |
| SM d18:0/16:0 | 19,142.55 | 10,433.59 | 52,340.65 | 85,042.84 | **0.014** | **0.000** | 173% |
| SM d18:1/17:1 | 491.95 | 303.25 | 1,422.02 | 2,098.66 | **0.006** | **0.000** | 189% |
| SM d18:1/17:0 | 3,208.37 | 1,806.82 | 9,836.29 | 15,034.51 | **0.006** | **0.000** | 207% |
| SM d18:1/18:1 | 18,640.25 | 9,808.52 | 43,836.62 | 65,728.79 | **0.016** | **0.001** | 135% |
| SM d18:1/18:0 | 40,892.78 | 20,053.25 | 90,535.08 | 135,225.95 | **0.021** | **0.002** | 121% |
| SM d18:0/18:0 | 6,790.95 | 4,093.27 | 10,526.87 | 14,699.17 | 0.116 | 0.452 | 55% |
| SM d18:1/19:1 | 447.89 | 234.09 | 1,460.18 | 2,219.10 | **0.004** | **0.000** | 226% |
| SM d18:1/19:0 | 1,787.55 | 822.65 | 4,662.20 | 6,747.23 | **0.007** | **0.000** | 161% |
| SM d18:1/20:1 | 8,343.49 | 4,449.21 | 24,634.38 | 36,978.63 | **0.006** | **0.000** | 195% |
| SM d18:1/20:0 | 25,222.21 | 11,121.24 | 62,633.45 | 95,752.40 | **0.014** | **0.000** | 148% |
| SM d18:1/21:1 | 1,046.96 | 466.41 | 2,833.69 | 3,938.79 | **0.004** | **0.000** | 171% |
| SM d18:1/21:0 | 5,055.41 | 2,199.41 | 13,027.86 | 19,640.68 | **0.011** | **0.000** | 158% |
| SM d18:1/22:1 | 31,914.57 | 14,523.51 | 88,831.88 | 133,223.32 | **0.007** | **0.000** | 178% |
| SM d18:1/22:0 | 52,701.82 | 23,465.98 | 129,288.92 | 189,096.71 | **0.011** | **0.000** | 145% |
| SM d18:1/23:1 | 11,568.81 | 4,957.16 | 35,011.67 | 50,500.29 | **0.004** | **0.000** | 203% |
| SM d18:1/23:0 | 18,495.56 | 8,310.51 | 46,935.02 | 65,107.29 | **0.006** | **0.000** | 154% |
| SM d18:1/24:1 | 106,811.38 | 40,160.09 | 263,648.05 | 356,332.79 | **0.006** | **0.000** | 147% |
| SM d18:1/24:0 | 44,159.97 | 18,958.22 | 108,004.11 | 149,031.73 | **0.007** | **0.000** | 145% |
| SM d18:0/24:0 | 4,760.18 | 2,080.05 | 10,957.67 | 14,970.47 | **0.009** | **0.000** | 130% |
| SM d18:1/25:1 | 2,455.32 | 955.44 | 6,022.50 | 7,390.92 | **0.003** | **0.000** | 145% |
| SM d18:1/25:0 | 1,984.13 | 790.49 | 5,076.28 | 6,684.72 | **0.004** | **0.000** | 156% |
| SM d18:0/25:0 | 1,065.26 | 393.11 | 3,199.24 | 4,379.92 | **0.002** | **0.000** | 200% |
| SM d18:1/26:1 | 688.89 | 294.16 | 1,769.72 | 2,118.06 | **0.002** | **0.000** | 157% |
| SM d18:1/26:0 | 331.54 | 179.75 | 917.12 | 955.55 | **0.000** | **0.000** | 177% |
| SM d18:0/26:0 | 131.90 | 63.94 | 420.75 | 433.64 | **0.000** | **0.000** | 219% |
| Cer d18:1/16:0 | 728.41 | 354.42 | 728.69 | 250.48 | 0.997 | 0.369 | 0% |
| Cer d18:1/18:0 | 411.52 | 368.97 | 372.52 | 216.29 | 0.549 | 0.876 | -9% |
| Cer d18:0/18:0 | 243.21 | 242.38 | 259.16 | 193.09 | 0.736 | 0.204 | 7% |
| Cer d18:1/20:0 | 363.41 | 428.94 | 271.26 | 163.99 | 0.188 | 0.442 | -25% |
| Cer d18:1/22:0 | 1,109.08 | 461.70 | 952.66 | 482.30 | 0.129 | **0.045** | -14% |
| Cer d18:1/24:1 | 2,579.23 | 1,670.39 | 1,871.32 | 911.86 | **0.016** | **0.009** | -27% |
| Cer d18:1/24:0 | 3,348.22 | 1,577.30 | 2,789.03 | 1,280.18 | 0.074 | **0.037** | -17% |
| Cer d18:0/24:1 | 211.93 | 188.27 | 212.47 | 160.36 | 0.988 | 0.447 | 0% |
| GluCer d18:1/16:0 | 743.11 | 288.65 | 769.57 | 264.47 | 0.658 | 0.557 | 4% |
| GluCer d18:0/16:0 | 27.51 | 9.87 | 26.66 | 8.26 | 0.666 | 0.853 | -3% |
| GluCer d18:1/18:0 | 98.66 | 35.95 | 107.11 | 42.88 | 0.326 | 0.471 | 9% |
| GluCer d18:0/18:0 | 7.15 | 3.44 | 7.91 | 3.75 | 0.334 | 0.548 | 11% |
| GluCer d18:1/20:0 | 79.67 | 29.19 | 97.02 | 31.86 | **0.010** | **0.012** | 22% |
| GluCer d18:0/20:0 | 6.16 | 2.88 | 6.77 | 3.86 | 0.408 | 0.598 | 10% |
| GluCer d18:1/22:0 | 673.39 | 265.07 | 782.15 | 306.00 | 0.082 | 0.079 | 16% |
| GluCer d18:0/22:0 | 17.83 | 8.13 | 18.71 | 7.42 | 0.603 | 0.500 | 5% |
| GluCer d18:1/24:1 | 911.17 | 286.78 | 1,004.74 | 306.84 | 0.148 | 0.135 | 10% |
| GluCer d18:0/24:1 | 876.34 | 374.31 | 995.37 | 407.05 | 0.162 | 0.128 | 14% |
| GluCer d18:1/24:0 | 25.54 | 10.59 | 28.49 | 11.46 | 0.219 | 0.145 | 12% |
| GluCer d18:0/24:0 | 20.89 | 8.47 | 22.80 | 10.31 | 0.350 | 0.397 | 9% |
| LacCer d18:1/16:0 | 3,241.04 | 973.12 | 3,270.18 | 986.25 | 0.891 | 1.000 | 1% |
| LacCer d18:0/16:0 | 78.43 | 21.02 | 76.41 | 24.58 | 0.683 | 0.487 | -3% |
| LacCer d18:1/18:0 | 100.36 | 33.07 | 101.51 | 64.52 | 0.918 | 0.560 | 1% |
| LacCer d18:1/20:0 | 42.51 | 17.91 | 40.27 | 23.77 | 0.624 | 0.280 | -5% |
| LacCer d18:1/22:0 | 171.49 | 62.64 | 185.52 | 105.15 | 0.457 | 0.966 | 8% |
| LacCer d18:1/24:1 | 625.25 | 231.64 | 670.53 | 239.75 | 0.376 | 0.402 | 7% |
| LacCer d18:1/24:0 | 249.44 | 90.21 | 283.84 | 145.54 | 0.194 | 0.427 | 14% |
| LacCer d18:0/24:1 | 16.56 | 8.50 | 18.02 | 6.79 | 0.382 | 0.175 | 9% |
| Gb3 d18:1/16:0 | 1,413.50 | 368.06 | 1,453.47 | 335.65 | 0.600 | 0.233 | 3% |
| Gb3 d18:1/18:0 | 155.48 | 52.51 | 173.19 | 54.54 | 0.129 | 0.134 | 11% |
| Gb3 d18:1/20:0 | 66.58 | 26.62 | 72.58 | 24.25 | 0.278 | 0.251 | 9% |
| Gb3 d18:1/22:0 | 207.11 | 75.27 | 224.80 | 74.66 | 0.277 | 0.312 | 9% |
| Gb3 d18:1/24:1 | 28.47 | 16.65 | 24.78 | 12.29 | 0.244 | 0.557 | -13% |
| Gb3 d18:1/24:0 | 70.76 | 27.77 | 88.15 | 30.02 | **0.007** | **0.002** | 25% |
| GM3 d18:1/16:0 | 3,465.41 | 967.06 | 3,231.81 | 767.81 | 0.217 | 0.333 | -7% |
| GM3 d18:0/16:0 | 744.81 | 204.69 | 664.61 | 172.39 | 0.052 | **0.049** | -11% |
| GM3 d18:1/18:1 | 386.54 | 116.55 | 359.59 | 125.75 | 0.306 | 0.213 | -7% |
| GM3 d18:1/18:0 | 1,531.53 | 398.74 | 1,315.67 | 402.21 | **0.014** | **0.004** | -14% |
| GM3 d18:0/18:0 | 577.43 | 144.19 | 491.82 | 114.25 | **0.003** | **0.003** | -15% |
| GM3 d18:1/20:1 | 200.15 | 64.44 | 183.06 | 69.09 | 0.240 | 0.126 | -9% |
| GM3 d18:1/20:0 | 704.50 | 243.43 | 583.90 | 194.15 | **0.013** | **0.018** | -17% |
| GM3 d18:0/20:0 | 205.34 | 79.19 | 154.17 | 51.27 | **0.001** | **0.001** | -25% |
| GM3 d18:1/22:1 | 888.84 | 260.81 | 812.60 | 253.38 | 0.173 | 0.152 | -9% |
| GM3 d18:1/22:0 | 1,980.84 | 679.39 | 1,696.60 | 555.08 | **0.036** | **0.024** | -14% |
| GM3 d18:0/22:0 | 574.80 | 225.67 | 421.35 | 174.14 | **0.001** | **0.000** | -27% |
| GM3 d18:1/24:1 | 2,471.21 | 795.58 | 2,384.20 | 759.93 | 0.605 | 0.691 | -4% |
| GM3 d18:1/24:0 | 1,837.56 | 583.45 | 1,768.44 | 527.24 | 0.566 | 0.500 | -4% |
| GM3 d18:0/24:0 | 522.42 | 137.40 | 422.84 | 119.75 | **0.001** | **0.001** | -19% |
| FFA | 10,867,348.82 | 882,985.02 | 10,460,387.14 | 918,832.05 | **0.039** | 0.052 | -4% |
| SM | 609,084.46 | 270,992.42 | 1,584,483.26 | 2,339,287.65 | **0.009** | **0.000** | 160% |
| Cer | 8,995.02 | 3,992.95 | 7,457.11 | 2,531.94 | **0.035** | **0.017** | -17% |
| GluCer | 3,487.42 | 1,227.31 | 3,867.30 | 1,317.24 | 0.171 | 0.126 | 11% |
| LacCer | 4,525.08 | 1,327.02 | 4,646.27 | 1,458.56 | 0.688 | 0.863 | 3% |
| GM3 | 16,091.36 | 4,163.69 | 14,490.66 | 3,701.17 | 0.063 | **0.048** | -10% |
| Gb3 | 1,941.90 | 493.99 | 2,036.97 | 473.61 | 0.365 | 0.120 | 5% |
| SM/Cer | 71.68 | 32.29 | 219.40 | 312.15 | **0.003** | **0.000** | 206% |

**Supplementary Table 5.** FPIR differences in individual lipid species

|  | FPIR<350 (n=32) |  | FPIR>350 (n=54) |  | P value |  |  |
| --- | --- | --- | --- | --- | --- | --- | --- |
| Sphingolipid | Average lipid concentration (nM) | Standard deviation | Average lipid concentration (nM) | Standard deviation | t.test | Man whitney u test | Percentage changed in FPIR |
| FFA22:6 | 30933.36 | 10802.90 | 26115.65 | 10581.45 | 0.764 | **0.009** | -16% |
| FFA22:5 | 37292.94 | 4346.40 | 33589.23 | 6014.57 | 0.151 | **0.000** | -10% |
| FFA22:4 | 46413.66 | 4107.59 | 45503.00 | 5288.10 | 0.305 | 0.201 | -2% |
| FFA20:5 | 38365.22 | 7441.80 | 37622.47 | 6354.22 | 0.737 | 0.636 | -2% |
| FFA20:4 | 72213.60 | 24411.97 | 71481.17 | 33693.18 | 0.106 | 0.093 | -1% |
| FFA20:3 | 73346.37 | 5041.58 | 71400.93 | 6898.52 | 0.109 | 0.124 | -3% |
| FFA18:3 | 21277.51 | 3718.62 | 20356.99 | 3794.93 | 0.745 | 0.088 | -4% |
| FFA18:2 | 897119.34 | 235720.26 | 731169.97 | 246480.70 | 0.806 | **0.000** | -18% |
| FFA18:1 | 397139.23 | 63927.86 | 338900.51 | 68191.46 | 0.677 | **0.000** | -15% |
| FFA18:0 | 2539078.10 | 174963.97 | 2429498.92 | 245429.58 | **0.005** | **0.037** | -4% |
| FFA16:1 | 907304.82 | 261234.17 | 867858.64 | 280367.87 | 0.471 | 0.348 | -4% |
| FFA16:0 | 2735074.86 | 189190.91 | 2610145.40 | 213170.04 | 0.750 | **0.001** | -5% |
| FFA14:0 | 231556.10 | 18307.13 | 228883.85 | 22111.71 | 0.124 | 0.761 | -1% |
| FA13:0 | 1392401.95 | 127368.27 | 1383750.44 | 200297.49 | **0.013** | 0.900 | -1% |
| FA12:0 | 1545982.45 | 151439.78 | 1581309.86 | 239513.03 | **0.022** | 0.163 | 2% |
| SM d18:1/16:1 | 28592.38 | 14527.93 | 45611.58 | 65693.88 | **0.019** | 0.567 | 60% |
| SM d18:1/16:0 | 220911.63 | 140418.89 | 424540.27 | 749081.64 | **0.017** | 0.344 | 92% |
| SM d18:0/16:0 | 22923.05 | 12621.92 | 43952.56 | 78344.84 | **0.017** | 0.475 | 92% |
| SM d18:1/17:1 | 609.42 | 405.39 | 1180.17 | 1943.09 | **0.012** | 0.235 | 94% |
| SM d18:1/17:0 | 4199.35 | 2892.12 | 8021.65 | 13902.99 | **0.017** | 0.304 | 91% |
| SM d18:1/18:1 | 21177.75 | 10539.15 | 37666.93 | 60571.47 | **0.011** | 0.555 | 78% |
| SM d18:1/18:0 | 46927.91 | 21311.30 | 77765.70 | 124578.97 | **0.016** | 0.971 | 66% |
| SM d18:0/18:0 | 7123.20 | 3194.99 | 9638.14 | 13621.68 | **0.023** | 0.284 | 35% |
| SM d18:1/19:1 | 561.22 | 403.34 | 1205.56 | 2049.53 | **0.012** | **0.043** | 115% |
| SM d18:1/19:0 | 2215.56 | 1298.20 | 3876.22 | 6227.21 | **0.018** | 0.491 | 75% |
| SM d18:1/20:1 | 10106.24 | 6944.83 | 20572.95 | 34096.73 | **0.012** | 0.051 | 104% |
| SM d18:1/20:0 | 29256.41 | 15003.03 | 53314.81 | 88037.81 | **0.017** | 0.280 | 82% |
| SM d18:1/21:1 | 1249.56 | 761.24 | 2382.75 | 3636.45 | **0.013** | 0.093 | 91% |
| SM d18:1/21:0 | 6145.86 | 3652.73 | 10905.28 | 18058.02 | **0.026** | 0.296 | 77% |
| SM d18:1/22:1 | 38397.92 | 25186.68 | 74449.64 | 122525.63 | **0.024** | **0.035** | 94% |
| SM d18:1/22:0 | 61899.83 | 35014.45 | 109655.45 | 173877.17 | **0.029** | 0.136 | 77% |
| SM d18:1/23:1 | 14967.10 | 11338.97 | 28656.60 | 46526.03 | **0.025** | 0.062 | 91% |
| SM d18:1/23:0 | 22748.17 | 14122.38 | 39148.39 | 60015.67 | **0.031** | 0.195 | 72% |
| SM d18:1/24:1 | 124436.27 | 64619.26 | 224159.84 | 328669.90 | **0.023** | 0.064 | 80% |
| SM d18:1/24:0 | 52357.79 | 29191.84 | 91323.15 | 137391.44 | **0.035** | 0.120 | 74% |
| SM d18:0/24:0 | 5631.16 | 3078.11 | 9293.85 | 13788.43 | **0.045** | 0.218 | 65% |
| SM d18:1/25:1 | 2932.18 | 1577.39 | 5079.33 | 6854.28 | **0.016** | 0.074 | 73% |
| SM d18:1/25:0 | 2412.81 | 1357.12 | 4249.62 | 6185.30 | **0.019** | 0.110 | 76% |
| SM d18:0/25:0 | 1406.07 | 927.20 | 2602.09 | 4060.15 | **0.023** | 0.068 | 85% |
| SM d18:1/26:1 | 848.64 | 474.83 | 1474.90 | 1974.77 | **0.025** | 0.095 | 74% |
| SM d18:1/26:0 | 450.29 | 362.28 | 738.31 | 892.52 | **0.037** | 0.054 | 64% |
| SM d18:0/26:0 | 196.95 | 173.68 | 328.71 | 406.61 | **0.041** | **0.046** | 67% |
| Cer d18:1/16:0 | 770.11 | 318.55 | 703.93 | 295.06 | 0.365 | 0.442 | -9% |
| Cer d18:1/18:0 | 402.53 | 372.72 | 385.07 | 250.03 | 0.728 | 1.000 | -4% |
| Cer d18:0/18:0 | 225.15 | 189.60 | 266.91 | 232.62 | 0.505 | 0.348 | 19% |
| Cer d18:1/20:0 | 376.14 | 475.69 | 280.78 | 178.31 | 0.068 | 0.543 | -25% |
| Cer d18:1/22:0 | 1228.05 | 575.67 | 911.13 | 362.82 | **0.024** | **0.006** | -26% |
| Cer d18:1/24:1 | 2712.80 | 1758.19 | 1923.27 | 995.91 | 0.051 | **0.007** | -29% |
| Cer d18:1/24:0 | 3687.60 | 1795.29 | 2691.46 | 1058.12 | **0.043** | **0.002** | -27% |
| Cer d18:0/24:1 | 224.58 | 194.17 | 204.87 | 161.51 | 0.239 | 0.782 | -9% |
| GluCer d18:1/16:0 | 834.39 | 295.06 | 710.58 | 254.47 | 0.113 | **0.033** | -15% |
| GluCer d18:0/16:0 | 27.16 | 9.33 | 27.02 | 8.95 | 0.718 | 0.865 | 0% |
| GluCer d18:1/18:0 | 108.75 | 35.64 | 99.57 | 41.78 | 0.875 | 0.109 | -8% |
| GluCer d18:0/18:0 | 7.27 | 3.66 | 7.70 | 3.59 | 0.334 | 0.506 | 6% |
| GluCer d18:1/20:0 | 92.66 | 33.03 | 86.11 | 30.83 | 0.612 | 0.353 | -7% |
| GluCer d18:0/20:0 | 6.42 | 3.19 | 6.50 | 3.56 | 0.793 | 0.964 | 1% |
| GluCer d18:1/22:0 | 762.16 | 302.54 | 709.41 | 283.78 | 0.350 | 0.427 | -7% |
| GluCer d18:0/22:0 | 17.55 | 7.97 | 18.71 | 7.65 | 0.989 | 0.458 | 7% |
| GluCer d18:1/24:1 | 983.05 | 312.80 | 944.82 | 292.86 | 0.718 | 0.491 | -4% |
| GluCer d18:0/24:1 | 1003.47 | 420.78 | 897.99 | 375.21 | 0.178 | 0.272 | -11% |
| GluCer d18:1/24:0 | 27.12 | 10.58 | 27.01 | 11.46 | 0.914 | 0.872 | 0% |
| GluCer d18:0/24:0 | 23.09 | 8.77 | 21.14 | 9.83 | 0.915 | 0.240 | -8% |
| LacCer d18:1/16:0 | 3512.77 | 975.36 | 3103.76 | 949.85 | 0.796 | **0.047** | -12% |
| LacCer d18:0/16:0 | 82.38 | 20.05 | 74.44 | 23.97 | 0.342 | 0.053 | -10% |
| LacCer d18:1/18:0 | 113.33 | 74.54 | 93.61 | 28.74 | **0.044** | 0.195 | -17% |
| LacCer d18:1/20:0 | 51.35 | 27.61 | 35.45 | 12.91 | **0.003** | **0.003** | -31% |
| LacCer d18:1/22:0 | 219.30 | 118.64 | 154.59 | 47.60 | **0.000** | **0.008** | -30% |
| LacCer d18:1/24:1 | 722.00 | 282.60 | 604.81 | 192.50 | **0.010** | 0.080 | -16% |
| LacCer d18:1/24:0 | 321.65 | 165.22 | 234.67 | 71.61 | **0.000** | **0.007** | -27% |
| LacCer d18:0/24:1 | 18.57 | 8.54 | 16.56 | 7.07 | 0.187 | 0.255 | -11% |
| Gb3 d18:1/16:0 | 1507.19 | 393.94 | 1390.55 | 317.68 | 0.489 | 0.057 | -8% |
| Gb3 d18:1/18:0 | 169.87 | 61.39 | 161.38 | 49.42 | 0.290 | 0.721 | -5% |
| Gb3 d18:1/20:0 | 70.80 | 27.65 | 68.97 | 24.33 | 0.485 | 0.796 | -3% |
| Gb3 d18:1/22:0 | 221.33 | 74.61 | 213.10 | 75.83 | 0.860 | 0.543 | -4% |
| Gb3 d18:1/24:1 | 27.09 | 16.33 | 26.28 | 13.64 | 0.191 | 0.886 | -3% |
| Gb3 d18:1/24:0 | 79.64 | 35.20 | 79.66 | 26.94 | 0.127 | 0.837 | 0% |
| GM3 d18:1/16:0 | 3513.20 | 950.06 | 3246.75 | 817.94 | 0.349 | 0.070 | -8% |
| GM3 d18:0/16:0 | 762.83 | 190.85 | 668.78 | 185.67 | 0.568 | **0.013** | -12% |
| GM3 d18:1/18:1 | 396.85 | 125.27 | 358.48 | 117.88 | 0.296 | 0.071 | -10% |
| GM3 d18:1/18:0 | 1535.46 | 415.03 | 1353.32 | 399.76 | 0.473 | **0.022** | -12% |
| GM3 d18:0/18:0 | 572.90 | 151.58 | 510.36 | 121.33 | 0.112 | 0.073 | -11% |
| GM3 d18:1/20:1 | 194.77 | 55.15 | 189.41 | 73.57 | 0.312 | 0.353 | -3% |
| GM3 d18:1/20:0 | 710.84 | 259.53 | 602.48 | 196.14 | 0.065 | 0.076 | -15% |
| GM3 d18:0/20:0 | 204.08 | 79.28 | 164.39 | 61.42 | **0.034** | **0.014** | -19% |
| GM3 d18:1/22:1 | 906.98 | 277.97 | 815.97 | 242.35 | 0.591 | 0.063 | -10% |
| GM3 d18:1/22:0 | 2038.86 | 729.71 | 1714.86 | 537.18 | 0.113 | **0.041** | -16% |
| GM3 d18:0/22:0 | 568.51 | 238.55 | 453.49 | 187.74 | 0.341 | **0.011** | -20% |
| GM3 d18:1/24:1 | 2582.12 | 800.58 | 2334.58 | 750.43 | 0.509 | 0.158 | -10% |
| GM3 d18:1/24:0 | 1967.27 | 574.02 | 1704.37 | 521.42 | 0.508 | **0.024** | -13% |
| GM3 d18:0/24:0 | 530.96 | 148.20 | 436.23 | 118.36 | 0.223 | **0.002** | -18% |
| FFA | 10965635.23 | 805437.08 | 10477506.62 | 940992.26 | 0.305 | **0.013** | -4% |
| SM | 730684.69 | 410181.86 | 1331794.46 | 2155670.38 | **0.019** | 0.198 | 82% |
| Cer | 9626.95 | 4404.06 | 7367.43 | 2283.34 | **0.030** | **0.003** | -23% |
| GluCer | 3893.08 | 1366.10 | 3556.56 | 1223.52 | 0.142 | 0.245 | -9% |
| LacCer | 5041.34 | 1538.19 | 4317.89 | 1229.60 | 0.101 | **0.029** | -14% |
| GM3 | 16485.62 | 4372.49 | 14553.45 | 3600.83 | 0.154 | **0.039** | -12% |
| Gb3 | 2075.91 | 549.72 | 1939.95 | 436.65 | 0.463 | 0.153 | -7% |
| SM/Cer | 78.81 | 35.76 | 187.82 | 288.88 | **0.005** | **0.002** | 138% |

**Supplementary Table 6.** Pearson’s relationship between lipids and clinical characteristics adjusted by age, sex, BMI, TG, TC, HDLc, LDLc, ALT and AST.

| lipid | GIR |  | GIR(age, sex) |  | GIR(age,sex,LDLc,HDLc,TC,TG) |  | GIR(age,sex,LDLc,HDLc,TC,TG,BMI) |  | GIR(age,sex,LDLc,HDLc,TC,TG,BMI,AST,ALT) |  |
| --- | --- | --- | --- | --- | --- | --- | --- | --- | --- | --- |
|  | r | p | adjusted r | p | adjusted r | p | adjusted r | p | adjusted r | p |
| FFA22:6 | 0.111 | 0.308 | **0.140** | **0.034** | **0.238** | **0.034** | **0.274** | **0.015** | **0.266** | **0.019** |
| FFA22:5 | -0.138 | 0.204 | -0.079 | 0.103 | 0.184 | 0.103 | **0.259** | **0.021** | **0.247** | **0.031** |
| FFA22:4 | **-.242\*** | **0.025** | -0.202 | 0.871 | -0.019 | 0.871 | 0.118 | 0.300 | 0.078 | 0.501 |
| FFA20:5 | 0.131 | 0.230 | 0.126 | 0.292 | 0.119 | 0.292 | **0.270** | **0.016** | **0.245** | **0.031** |
| FFA20:4 | **.304\*\*** | **0.004** | **0.304** | **0.001** | **0.360** | **0.001** | **0.455** | **0.000** | **0.428** | **0.000** |
| FFA20:3 | -0.066 | 0.544 | -0.024 | 0.524 | 0.072 | 0.524 | 0.162 | 0.154 | 0.138 | 0.230 |
| FFA18:3 | **-.287\*\*** | **0.007** | -0.242 | 0.717 | -0.041 | 0.717 | 0.062 | 0.588 | 0.045 | 0.695 |
| FFA18:2 | -0.186 | 0.087 | -0.123 | 0.544 | 0.069 | 0.544 | 0.190 | 0.093 | 0.164 | 0.154 |
| FFA18:1 | **-.384\*\*** | **0.000** | -0.340 | 0.370 | -0.102 | 0.370 | 0.008 | 0.942 | 0.002 | 0.987 |
| FFA18:0 | **-.498\*\*** | **0.000** | **-0.495** | **0.009** | **-0.291** | **0.009** | -0.176 | 0.120 | -0.185 | 0.108 |
| FFA16:1 | **-.555\*\*** | **0.000** | **-0.548** | **0.000** | **-0.396** | **0.000** | -0.214 | 0.058 | **-0.243** | **0.033** |
| FFA16:0 | **-.294\*\*** | **0.006** | -0.283 | 0.506 | -0.076 | 0.506 | 0.028 | 0.807 | 0.007 | 0.952 |
| FFA14:0 | -0.207 | 0.056 | -0.182 | 0.656 | -0.051 | 0.656 | 0.050 | 0.660 | 0.019 | 0.869 |
| FA13:0 | 0.151 | 0.166 | 0.172 | 0.294 | 0.119 | 0.294 | 0.093 | 0.414 | 0.077 | 0.504 |
| FA12:0 | **.239\*** | **0.027** | 0.250 | 0.270 | 0.125 | 0.270 | 0.066 | 0.564 | 0.037 | 0.748 |
| SM d18:1/16:1 | **.441\*\*** | **0.000** | **0.420** | **0.000** | **0.403** | **0.000** | **0.313** | **0.005** | **0.317** | **0.005** |
| SM d18:1/16:0 | **.468\*\*** | **0.000** | **0.449** | **0.000** | **0.422** | **0.000** | **0.319** | **0.004** | **0.323** | **0.004** |
| SM d18:0/16:0 | **.442\*\*** | **0.000** | **0.422** | **0.000** | **0.405** | **0.000** | **0.304** | **0.007** | **0.308** | **0.006** |
| SM d18:1/17:1 | **.483\*\*** | **0.000** | **0.466** | **0.000** | **0.430** | **0.000** | **0.325** | **0.003** | **0.330** | **0.003** |
| SM d18:1/17:0 | **.482\*\*** | **0.000** | **0.466** | **0.000** | **0.432** | **0.000** | **0.322** | **0.004** | **0.328** | **0.004** |
| SM d18:1/18:1 | **.435\*\*** | **0.000** | **0.416** | **0.000** | **0.400** | **0.000** | **0.293** | **0.009** | **0.298** | **0.008** |
| SM d18:1/18:0 | **.424\*\*** | **0.000** | **0.405** | **0.000** | **0.392** | **0.000** | **0.287** | **0.010** | **0.294** | **0.009** |
| SM d18:0/18:0 | **.323\*\*** | **0.002** | **0.300** | **0.003** | **0.328** | **0.003** | **0.239** | **0.034** | **0.250** | **0.028** |
| SM d18:1/19:1 | **.495\*\*** | **0.000** | **0.477** | **0.000** | **0.439** | **0.000** | **0.330** | **0.003** | **0.333** | **0.003** |
| SM d18:1/19:0 | **.469\*\*** | **0.000** | **0.456** | **0.000** | **0.424** | **0.000** | **0.317** | **0.004** | **0.325** | **0.004** |
| SM d18:1/20:1 | **.484\*\*** | **0.000** | **0.466** | **0.000** | **0.432** | **0.000** | **0.326** | **0.003** | **0.330** | **0.003** |
| SM d18:1/20:0 | **.444\*\*** | **0.000** | **0.428** | **0.000** | **0.410** | **0.000** | **0.305** | **0.006** | **0.311** | **0.006** |
| SM d18:1/21:1 | **.494\*\*** | **0.000** | **0.477** | **0.000** | **0.449** | **0.000** | **0.345** | **0.002** | **0.347** | **0.002** |
| SM d18:1/21:0 | **.463\*\*** | **0.000** | **0.449** | **0.000** | **0.427** | **0.000** | **0.328** | **0.003** | **0.335** | **0.003** |
| SM d18:1/22:1 | **.479\*\*** | **0.000** | **0.461** | **0.000** | **0.431** | **0.000** | **0.333** | **0.003** | **0.337** | **0.003** |
| SM d18:1/22:0 | **.463\*\*** | **0.000** | **0.445** | **0.000** | **0.424** | **0.000** | **0.329** | **0.003** | **0.332** | **0.003** |
| SM d18:1/23:1 | **.512\*\*** | **0.000** | **0.495** | **0.000** | **0.462** | **0.000** | **0.359** | **0.001** | **0.363** | **0.001** |
| SM d18:1/23:0 | **.492\*\*** | **0.000** | **0.477** | **0.000** | **0.450** | **0.000** | **0.355** | **0.001** | **0.359** | **0.001** |
| SM d18:1/24:1 | **.473\*\*** | **0.000** | **0.458** | **0.000** | **0.438** | **0.000** | **0.338** | **0.002** | **0.339** | **0.003** |
| SM d18:1/24:0 | **.474\*\*** | **0.000** | **0.458** | **0.000** | **0.431** | **0.000** | **0.340** | **0.002** | **0.345** | **0.002** |
| SM d18:0/24:0 | **.464\*\*** | **0.000** | **0.449** | **0.000** | **0.428** | **0.000** | **0.340** | **0.002** | **0.346** | **0.002** |
| SM d18:1/25:1 | **.507\*\*** | **0.000** | **0.495** | **0.000** | **0.446** | **0.000** | **0.343** | **0.002** | **0.351** | **0.002** |
| SM d18:1/25:0 | **.506\*\*** | **0.000** | **0.491** | **0.000** | **0.447** | **0.000** | **0.343** | **0.002** | **0.349** | **0.002** |
| SM d18:0/25:0 | **.519\*\*** | **0.000** | **0.503** | **0.000** | **0.460** | **0.000** | **0.353** | **0.001** | **0.354** | **0.002** |
| SM d18:1/26:1 | **.514\*\*** | **0.000** | **0.502** | **0.000** | **0.464** | **0.000** | **0.368** | **0.001** | **0.368** | **0.001** |
| SM d18:1/26:0 | **.587\*\*** | **0.000** | **0.571** | **0.000** | **0.507** | **0.000** | **0.431** | **0.000** | **0.429** | **0.000** |
| SM d18:0/26:0 | **.642\*\*** | **0.000** | **0.628** | **0.000** | **0.552** | **0.000** | **0.463** | **0.000** | **0.463** | **0.000** |
| Cer d18:1/16:0 | 0.041 | 0.707 | 0.043 | 0.996 | -0.001 | 0.996 | 0.004 | 0.972 | -0.001 | 0.994 |
| Cer d18:1/18:0 | -0.062 | 0.571 | -0.082 | 0.411 | -0.093 | 0.411 | 0.111 | 0.331 | 0.073 | 0.526 |
| Cer d18:0/18:0 | 0.016 | 0.886 | 0.008 | 0.735 | 0.038 | 0.735 | 0.028 | 0.809 | 0.024 | 0.837 |
| Cer d18:1/20:0 | -0.145 | 0.184 | -0.147 | 0.653 | -0.051 | 0.653 | 0.141 | 0.216 | 0.128 | 0.269 |
| Cer d18:1/22:0 | -0.195 | 0.071 | -0.194 | 0.998 | 0.000 | 0.998 | -0.033 | 0.771 | -0.022 | 0.853 |
| Cer d18:1/24:1 | **-.263\*** | **0.014** | -0.247 | 0.370 | -0.102 | 0.370 | 0.021 | 0.853 | 0.003 | 0.982 |
| Cer d18:1/24:0 | -0.186 | 0.087 | -0.150 | 0.614 | 0.057 | 0.614 | -0.005 | 0.967 | 0.018 | 0.875 |
| Cer d18:0/24:1 | 0.028 | 0.796 | 0.034 | 0.863 | 0.020 | 0.863 | 0.086 | 0.450 | 0.065 | 0.576 |
| GluCer d18:1/16:0 | 0.056 | 0.608 | 0.030 | 0.126 | 0.172 | 0.126 | 0.121 | 0.287 | 0.145 | 0.209 |
| GluCer d18:0/16:0 | 0.009 | 0.936 | -0.005 | 0.404 | 0.095 | 0.404 | 0.113 | 0.322 | 0.112 | 0.332 |
| GluCer d18:1/18:0 | 0.095 | 0.384 | 0.087 | 0.164 | 0.157 | 0.164 | 0.005 | 0.967 | 0.018 | 0.877 |
| GluCer d18:0/18:0 | 0.105 | 0.337 | 0.064 | 0.073 | 0.202 | 0.073 | 0.037 | 0.744 | 0.086 | 0.458 |
| GluCer d18:1/20:0 | **.262\*** | **0.015** | **0.253** | **0.006** | **0.302** | **0.006** | 0.164 | 0.150 | 0.169 | 0.141 |
| GluCer d18:0/20:0 | 0.057 | 0.604 | 0.046 | 0.835 | 0.024 | 0.835 | 0.068 | 0.554 | 0.043 | 0.714 |
| GluCer d18:1/22:0 | **.233\*** | **0.031** | **0.219** | **0.001** | **0.365** | **0.001** | **0.241** | **0.032** | **0.237** | **0.038** |
| GluCer d18:0/22:0 | 0.145 | 0.182 | 0.107 | 0.143 | 0.165 | 0.143 | 0.101 | 0.375 | 0.072 | 0.535 |
| GluCer d18:1/24:1 | 0.186 | 0.086 | **0.153** | **0.012** | **0.280** | **0.012** | 0.211 | 0.063 | 0.196 | 0.087 |
| GluCer d18:0/24:1 | 0.169 | 0.121 | **0.163** | **0.006** | **0.307** | **0.006** | **0.226** | **0.045** | **0.229** | **0.045** |
| GluCer d18:1/24:0 | 0.195 | 0.071 | **0.170** | **0.004** | **0.321** | **0.004** | **0.242** | **0.031** | **0.236** | **0.039** |
| GluCer d18:0/24:0 | 0.115 | 0.293 | 0.081 | 0.130 | 0.171 | 0.130 | 0.197 | 0.081 | 0.190 | 0.098 |
| LacCer d18:1/16:0 | 0.046 | 0.675 | 0.071 | 0.230 | 0.136 | 0.230 | 0.112 | 0.325 | 0.130 | 0.259 |
| LacCer d18:0/16:0 | -0.012 | 0.909 | 0.001 | 0.731 | -0.039 | 0.731 | 0.013 | 0.909 | 0.025 | 0.829 |
| LacCer d18:1/18:0 | 0.185 | 0.089 | 0.198 | 0.385 | 0.099 | 0.385 | 0.101 | 0.378 | 0.125 | 0.278 |
| LacCer d18:1/20:0 | -0.023 | 0.834 | -0.033 | 0.649 | -0.052 | 0.649 | -0.073 | 0.524 | -0.035 | 0.759 |
| LacCer d18:1/22:0 | 0.143 | 0.190 | 0.163 | 0.110 | 0.180 | 0.110 | 0.141 | 0.215 | 0.163 | 0.156 |
| LacCer d18:1/24:1 | 0.180 | 0.097 | 0.183 | 0.328 | 0.111 | 0.328 | 0.061 | 0.596 | 0.087 | 0.450 |
| LacCer d18:1/24:0 | **.239\*** | **0.027** | 0.259 | 0.115 | 0.178 | 0.115 | 0.151 | 0.185 | 0.200 | 0.081 |
| LacCer d18:0/24:1 | 0.094 | 0.387 | 0.129 | 0.283 | 0.121 | 0.283 | -0.015 | 0.896 | -0.021 | 0.855 |
| Gb3 d18:1/16:0 | 0.193 | 0.075 | **0.193** | **0.036** | **0.235** | **0.036** | **0.277** | **0.013** | **0.256** | **0.025** |
| Gb3 d18:1/18:0 | **.267\*** | **0.013** | **0.288** | **0.019** | **0.261** | **0.019** | 0.122 | 0.283 | 0.070 | 0.542 |
| Gb3 d18:1/20:0 | 0.164 | 0.131 | 0.164 | 0.358 | 0.104 | 0.358 | 0.045 | 0.695 | 0.051 | 0.657 |
| Gb3 d18:1/22:0 | **.285\*\*** | **0.008** | **0.290** | **0.045** | **0.225** | **0.045** | **0.252** | **0.025** | 0.219 | 0.055 |
| Gb3 d18:1/24:1 | 0.033 | 0.766 | 0.004 | 0.858 | 0.020 | 0.858 | 0.061 | 0.592 | 0.041 | 0.725 |
| Gb3 d18:1/24:0 | **.351\*\*** | **0.001** | **0.363** | **0.030** | **0.243** | **0.030** | **0.270** | **0.016** | **0.264** | **0.020** |
| GM3 d18:1/16:0 | -0.157 | 0.148 | -0.148 | 0.986 | -0.002 | 0.986 | -0.054 | 0.634 | -0.001 | 0.996 |
| GM3 d18:0/16:0 | **-.244\*** | **0.023** | -0.209 | 0.580 | -0.063 | 0.580 | -0.076 | 0.507 | -0.022 | 0.849 |
| GM3 d18:1/18:1 | -0.120 | 0.272 | -0.105 | 0.224 | -0.138 | 0.224 | -0.189 | 0.095 | -0.166 | 0.150 |
| GM3 d18:1/18:0 | **-.312\*\*** | **0.003** | **-0.280** | **0.027** | **-0.247** | **0.027** | **-0.284** | **0.011** | **-0.243** | **0.033** |
| GM3 d18:0/18:0 | **-.422\*\*** | **0.000** | **-0.397** | **0.002** | **-0.345** | **0.002** | **-0.344** | **0.002** | **-0.290** | **0.011** |
| GM3 d18:1/20:1 | **-.247\*** | **0.022** | -0.228 | 0.104 | -0.183 | 0.104 | -0.201 | 0.076 | -0.210 | 0.066 |
| GM3 d18:1/20:0 | **-.431\*\*** | **0.000** | **-0.393** | **0.017** | **-0.266** | **0.017** | **-0.295** | **0.008** | **-0.235** | **0.039** |
| GM3 d18:0/20:0 | **-.507\*\*** | **0.000** | **-0.490** | **0.000** | **-0.383** | **0.000** | **-0.350** | **0.002** | **-0.306** | **0.007** |
| GM3 d18:1/22:1 | **-.280\*\*** | **0.009** | -0.261 | 0.334 | -0.109 | 0.334 | -0.117 | 0.304 | -0.064 | 0.579 |
| GM3 d18:1/22:0 | **-.347\*\*** | **0.001** | -0.316 | 0.369 | -0.102 | 0.369 | -0.132 | 0.246 | -0.070 | 0.546 |
| GM3 d18:0/22:0 | **-.471\*\*** | **0.000** | -0.446 | 0.072 | -0.203 | 0.072 | **-0.235** | **0.037** | -0.172 | 0.135 |
| GM3 d18:1/24:1 | -0.180 | 0.097 | -0.150 | 0.542 | -0.069 | 0.542 | -0.199 | 0.079 | -0.140 | 0.223 |
| GM3 d18:1/24:0 | -0.147 | 0.178 | -0.106 | 0.942 | -0.008 | 0.942 | -0.118 | 0.299 | -0.049 | 0.670 |
| GM3 d18:0/24:0 | **-.400\*\*** | **0.000** | -0.377 | 0.111 | -0.179 | 0.111 | -0.165 | 0.146 | -0.115 | 0.317 |
| FFA | **-.350\*\*** | **0.001** | -0.316 | 0.284 | -0.121 | 0.284 | 0.021 | 0.855 | -0.015 | 0.894 |
| SM | **.467\*\*** | **0.000** | **0.449** | **0.000** | **0.425** | **0.000** | **0.324** | **0.004** | **0.328** | **0.004** |
| Cer | **-.227\*** | **0.036** | -0.205 | 0.786 | -0.031 | 0.786 | 0.039 | 0.735 | 0.035 | 0.765 |
| GluCer | 0.173 | 0.111 | **0.153** | **0.006** | **0.304** | **0.006** | 0.213 | 0.060 | 0.214 | 0.061 |
| LacCer | 0.099 | 0.362 | 0.122 | 0.201 | 0.144 | 0.201 | 0.114 | 0.316 | 0.139 | 0.228 |
| GM3 | **-.301\*\*** | **0.005** | -0.270 | 0.252 | -0.130 | 0.252 | -0.200 | 0.078 | -0.134 | 0.247 |
| Gb3 | **.245\*** | **0.023** | **0.248** | **0.016** | **0.268** | **0.016** | **0.287** | **0.010** | **0.260** | **0.022** |
| SM/Cer | **.498\*\*** | **0.000** | **0.478** | **0.000** | **0.434** | **0.000** | **0.318** | **0.004** | **0.321** | **0.004** |

**Supplementary Table 7.**
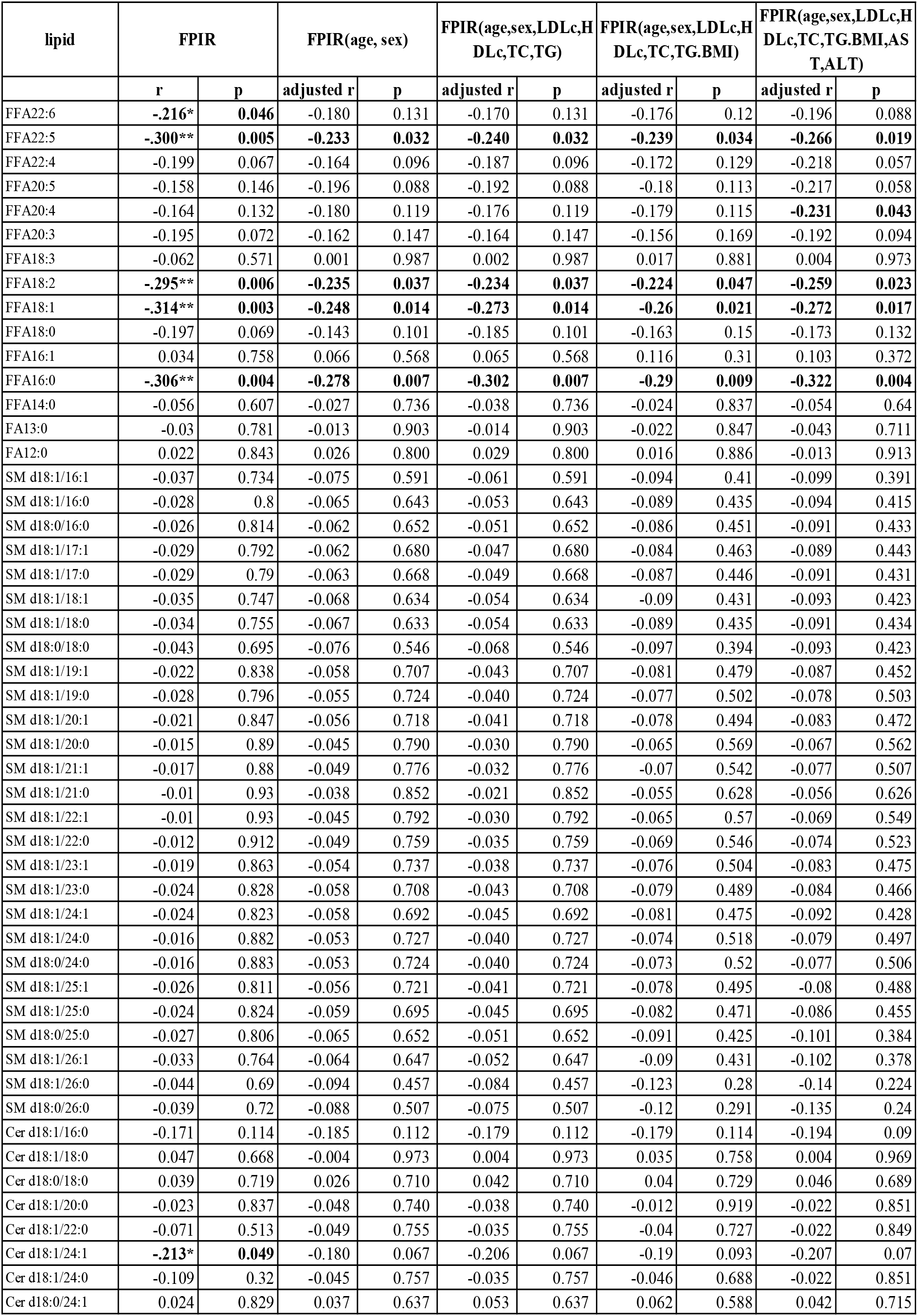
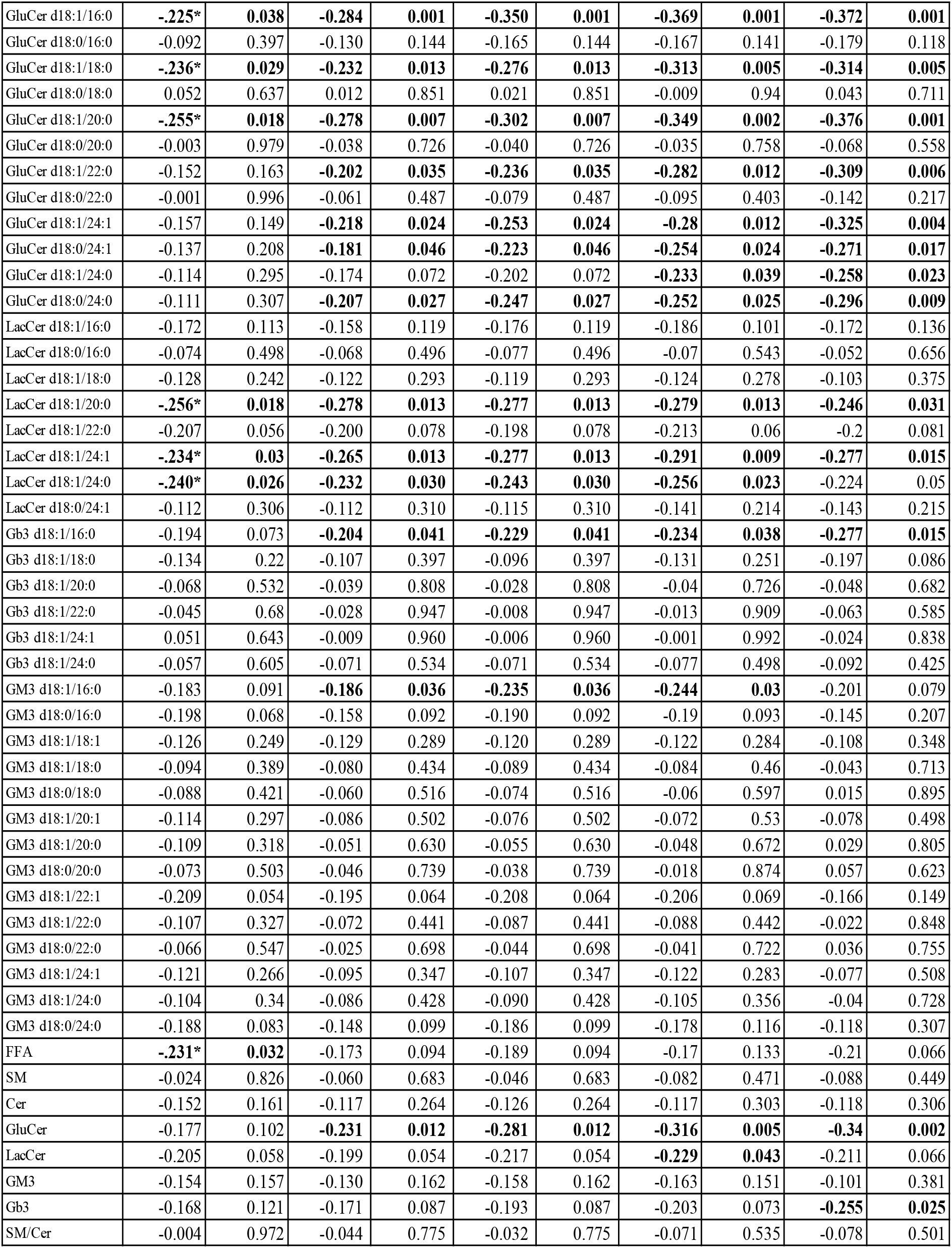
ROC curve for insulin sensitivity.

